# The Species Awareness Index (SAI): a Wikipedia-derived conservation culturomics metric for public biodiversity awareness

**DOI:** 10.1101/2020.08.17.254177

**Authors:** Joseph W Millard, Richard D Gregory, Kate Jones, Robin Freeman

## Abstract

Threats to global biodiversity are well-known, but slowing currents rates of biodiversity loss remains an ongoing challenge. The Aichi Targets set out 20 goals on which the international community should act to alleviate biodiversity decline, one of which (Target 1) aimed to raise public awareness of the importance of biodiversity. Whilst conventional indicators for Target 1 are of low spatial and temporal coverage, conservation culturomics has demonstrated how biodiversity awareness can be quantified at the global scale. Following the Living Planet Index methodology, here we introduced the Species Awareness Index (SAI), an index of changing species awareness on Wikipedia. We calculated this index at the page level for 41,197 IUCN species across 10 Wikipedia languages, incorporating over 2 billion views. Bootstrapped indices for the page level SAI show that overall awareness of biodiversity is marginally increasing, although there are differences among taxonomic classes and languages. Among taxonomic classes, overall awareness of reptiles is increasing fastest, and amphibians slowest. Among languages, overall species awareness for the Japanese Wikipedias is increasing fastest, and the Chinese and German Wikipedias slowest. Although awareness of species on Wikipedia as a whole is increasing, and is significantly higher in traded species, over the period 2016-2020 change in interest appears not to be strongly related to the trade of species or animal pollinators. As a data source for public biodiversity awareness Wikipedia could be integrated into the Biodiversity Engagement Indicator, thereby incorporating a more direct link to biodiversity itself.

## Introduction

Threats to global biodiversity are well-known, but slowing currents rates of biodiversity loss remains an ongoing challenge (IPBES, 2019; Mace et al., 2018). One problem is the requirement for transformational behavioural and economic change (IPBES, 2019), and the difficulty in leveraging this change at the global level across states (IPBES, 2019). The Strategic Plan for Biodiversity 2011–2020, underpinned by the Aichi Targets, represents an effort to guide these changes (UNEP CBD, 2010). Specifically, the Aichi Targets set out 20 goals on which the international community should act to alleviate biodiversity decline (UNEP CBD, 2010). Three of the Aichi Targets are well-evidenced (6, 9, and 11), four have intermediate sufficiency (4, 7, 12, and 14), ten are insufficient (1, 5, 8, 10, 13, 16, 17-20), and 3 have no indicators (Mcowen et al., 2016). Concerned with public awareness of biodiversity, Aichi Target 1 states that by 2020 the public should be aware of the value of biodiversity. Conventional indicators for Target 1 (i.e. the Biodiversity Barometer, UEBT 2019)) are of low spatial and temporal coverage (Leadley, 2013; Mcowen et al., 2016), and do not incorporate awareness of biodiversity itself (i.e. species). Without robust metrics capturing evidence towards Target 1, understanding whether this target has been met is hard.

Conservation culturomics has emerged as a field concerned broadly with digitised data and human nature interactions (Sherren et al., 2017; Ladle et al 2016). Quantifying public awareness of biodiversity is an area of active interest. Using data sources such as Twitter, Facebook, Flickr, Wikipedia, and Google Trends, a number of researchers have shown how online data can be used to better understand how the public perceives biodiversity and environmentalism (Roberge 2014; Papworth et al 2015; Tenkanen et al 2017; Roll et al 2016; Mccallum & Bury 2013). More recently, research has explored how online data sources can be combined to build a single indicator of biodiversity awareness (Cooper et al 2019). For example, Cooper et al. (2019) examined frequencies of biodiversity keywords across social media, online newspapers, and internet searches, reasoning that relative frequencies reflect public awareness of conservation issues. A significant step forward in applying culturomic approaches to the development of indicators, Cooper et al. (2019) provided a global framework for future research. Given Cooper et al (2019) focuses on conservation issues, one potential improvement could be to incorporate changing awareness of biodiversity itself.

Wikipedia page views represent a powerful data source for quantifying change in public awareness of biodiversity itself. Page views have previously been applied to quantify reptile public interest (Roll et al., 2016) and species phenology (Mittermeier et al., 2019). In the context of awareness, Wikipedia is valuable in that pages are linked explicitly to biodiversity across scales. Pages on Wikipedia exist for taxa at multiple taxonomic levels, Red List statuses, and ecological systems, with an unambiguous link between the taxon and page identity (Mittermeier et al., 2019). Previous research has shown Wikipedia can reveal changes in public awareness in response to natural history documentaries, demonstrating that the data source could be informative of long-term changes in awareness (Fernández-Bellon & Kane, 2019). Moreover, since species characteristics provide a mechanistic link to ecosystem services, change in awareness for a particular species on Wikipedia could be used as a proxy for awareness of its contribution. For example, increasing awareness for species which contribute significantly to pollination or provisioning services (i.e food and raw materials) could indicate greater public awareness of biodiversity importance. For pollination specifically, such changes in awareness are particularly important, given the global economic importance and reported declines of animal pollinators (Lautenbach et al. 2012; Powney et al. 2019; Hallmann et al. 2017; IPBES 2017). Although using Wikipedia for quantifying awareness is not without its limitations and caveats (see Discussion), it provides the basis for a useful additional indicator.

An awareness metric using Wikipedia page views could be thought of as analogous to The Living Planet Index (LPI). The LPI represents an aggregation of vertebrate population trends (Loh et al., 2005; Collen, 2009; McRae, 2017), showing an average rate of change for multiple species populations. Treating species page-views as a population size, the LPI methodology could be similarly applied to Wikipedia to derive a rate of change for species awareness. Multiple studies have used page views or search trends to infer change in awareness of specific species (Fink et al 2020; Fukano et al 2020; Lenda et al 2020; Sorian-Redondo et al 2017; Mittermeier et al 2019; Verissimo et al 2020), but as far as we know no studies have calculated such an aggregated index for overall awareness. Here, we introduce and evaluate an approach based on the frequency of views for IUCN species on Wikipedia, naming it the Species Awareness Index (SAI). We then explore how this metric varies for a series of groups, aiming to assess whether awareness of biodiversity has changed. Specifically, we explore biodiversity pooled, 6 distinct taxonomic classes (reptiles, ray-finned fishes, mammals, birds, insects, and amphibians), and each taxonomic class in each of 10 languages (Arabic, Chinese, English, French, German, Italian, Japanese, Portuguese, Russian, Spanish). We then predict rate of change in the page level SAI as a function of taxonomic class, Wikipedia language, trade status, and pollination contribution, using a pollinator dataset derived from the academic literature through named-entity recognition (Millard, et al., 2020). We conclude by discussing the limitations of the SAI, suggesting potential avenues for future research, and demonstrating how the SAI might be combined with other approaches for a more holistic understanding of changing biodiversity awareness.

## Methods

### Wikipedia data

We used the Wikipedia pageview API^1^, and software written in Python, to download daily user views of IUCN species for the period 1^st^ July 2015 – 30^1st^ March 2020 (downloaded on the 16^th^-21^st^ April 2020). We downloaded views for all IUCN species with Wikipedia pages in the taxonomic groups reptiles, ray-finned fishes (Actinopterygii), mammals, birds, insects, and amphibians, from 10 Wikipedia language projects (Arabic, Chinese, English, French, German, Italian, Japanese, Portuguese, Russian, Spanish). Each individual Wikipedia page for a particular species we henceforth refer to as a “species page”, distinguishing from our use of the term “species” to refer to a particular species among languages. Through the Wikipedia API, Onezoom (Rosindell & Wong, 2020) provided a reference for mapping between a species’ IUCN ID, Wikidata ID, and the main Wikipedia page name for each species in each language ^2^. Downloading views from only the main page name of each species excludes redirect views, controlling for potential variation caused by the URL used to reach a page. For each species page we retrieved only user views (i.e. views for which the visitor to that page was recorded as human, excluding automated views from bots). As in Mittermeier et al (2019), we were not able to retrieve views from before 1st July 2015, since views from before this date are not archived by Wikipedia at the pageview API. Future iterations of the SAI should strive towards reconciling views archived elsewhere across formats (see Discussion).

For each species page returned, we calculated the daily average views for each month, and then kept only those species pages for which the series was represented for all months (see Supplementary Information Figure S1 for the number of complete series). We used daily average views rather than total views since the Wikipedia pageview API does not always return views for all days in a given month.

To account for the overall change in Wikipedia’s popularity and use, we also downloaded the daily user views for a random set of 11000 pages in each language, using the Wikipedia Random API^3^ to request random pages. We then aggregated these views in the same manner as the daily average species views, and again kept only pages represented across the whole time series. From this random set of views, we then removed any page also appearing in the set of species pages for that language. We initially sampled 11,000 pages to maximise the number of remaining pages after removing incomplete series and species pages.

### Pollinator and wildlife trade datasets

To explore how species awareness varied with pollination contribution, we built a list of animal pollinators with an approach combining text-analysis and manual inspection of the pollination literature (see Supplementary Information for a detailed methodology). We also used the list of traded vertebrate species released in Scheffers et al (2019) and FAO fisheries statistics (FAO, 2020) to compile a dataset of traded mammals, birds, squamate reptiles, and harvested ray-finned fish. We then retrieved the Wikidata ID for each of these traded species using the Wikipedia API, which we merged onto each species page. In the following paper we henceforth refer to any species that pollinates as providing a “pollination contribution”, and any species in either Schefers et al (2019) or the FAO statistics as “traded”.

### Calculating absolute awareness of biodiversity

Before calculating the SAI we briefly explored absolute awareness of biodiversity among taxonomic classes, pollination contribution, and trade status. We defined “absolute awareness” as the total views for a species page on Wikipedia in the period 1^st^ July 2015 – 30^1st^ March 2020. We merged total views for each species page with the taxonomic class,

trade status, and pollination contribution of that species, and then built two generalised linear mixed-effects model: 1) modelling log_10_ total article views as a function of taxonomic class, trade status (Y/N), the interaction of class and trade, and a random effect for language; and 2) modelling log_10_ total article views as a function of taxonomic class, pollination contribution (Y/N), the interaction of class and pollination, and a random effect for language. Rather than attempting to find the most parsimonious model, we present full model predicted values, with AIC values for these and a set of candidate null models included in the Supplementary Information (Tables S6 and S7). In the Supplementary Information (Figures S11 and S12) we also present boxplots for the distribution of total views among taxa for each language.

### Deriving the Species Awareness Index (SAI)

The Species Awareness Index (SAI) is a new measurement of change in species awareness, calculated at the species page level from the rate of change in daily average Wikipedia views per month. Since the SAI measures the rate of change in views within a species page, species are weighted equally irrespective of their popularity, meaning highly viewed species do not dominate the SAI. In the remainder of this paper we use the term ‘SAI’ or ‘Species Awareness Index’ to refer to the overall change in awareness for a given species page, species, or group of species on Wikipedia. Specifically, we use the term “species page SAI” to refer to rate of change at the page level, the term “species SAI” to refer to the average of all species page SAIs for a unique species among languages, and “overall SAI” to refer to a bootstrapped group of species SAIs (see Figure 1). We also use the term “average monthly rate of change in the species page SAI” to refer to the average rate of change for a single species page across a given time period. All of the above are distinct from absolute interest in a given species or group of species (i.e. the total Wikipedia views over the whole time series).

**Figure 1.**
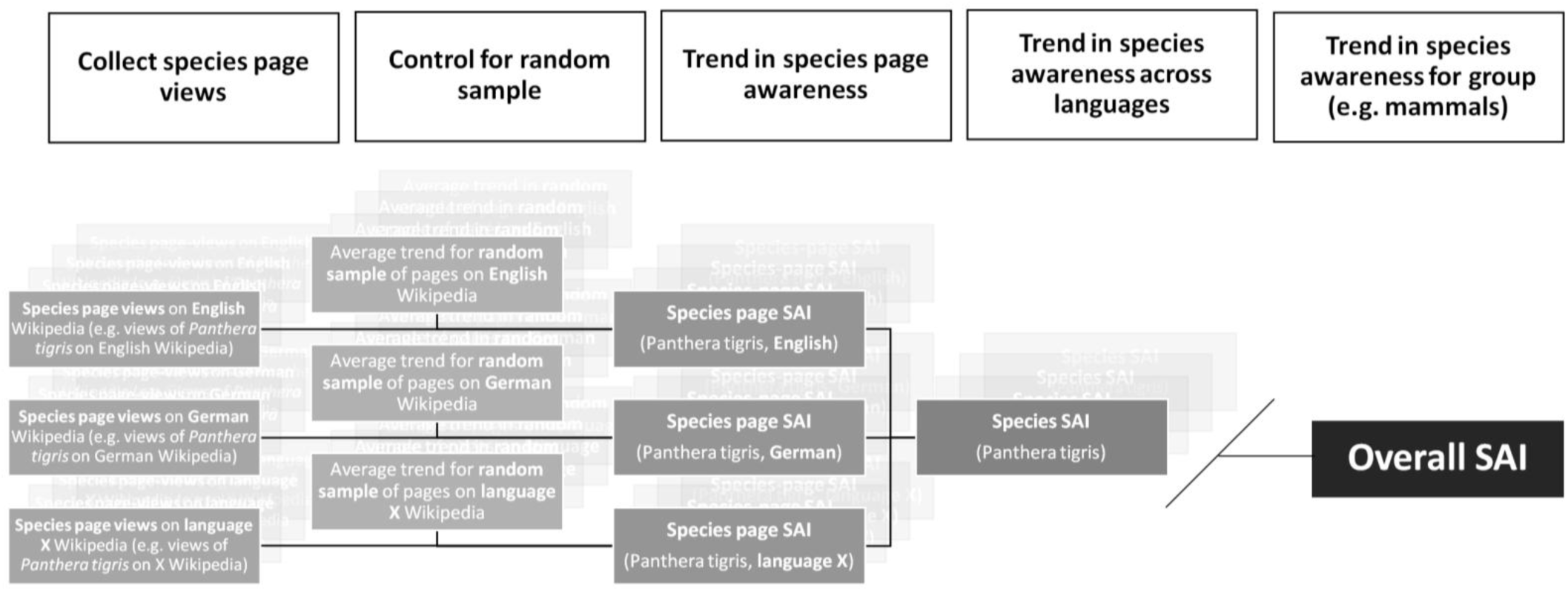
A schematic describing how the species page, species, and overall SAI were derived using Wikipedia views. The *species page SAI* represents the random adjusted trend for a given species in a given language, the *species SAI* is the average of species page SAIs for a single species across languages, and the *overall SAI* is a group of bootstrapped species SAIs.

To construct the species page SAI, we used the R package “rlpi” to calculate an index of change over time for each species in 6 taxonomic groups (amphibians, birds, insects, mammals, ray-finned fish, and reptiles) on 10 Wikipedia languages (Arabic, Chinese, English, French, German, Italian, Japanese, Portuguese, Russian, Spanish). The “rlpi” package applies a generalised additive model (GAM) to smooth the daily average species page view trends, using *k* = *N*/2 for the degrees of freedom parameter, following (Collen, 2009). In ‘rlpi’, these smoothed values are then used to calculate a rate of change in views for a species page article, as

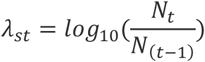

where *λs* = the rate of change in a species page, *N* = the smoothed number of daily average page views per month, and *t* = month.

To account for the overall change in popularity of Wikipedia itself over the same time period, we adjusted the rate of change for each species page using the rate of change in a random set of complete series Wikipedia pages (see Supplementary Information for the number of complete series, Figures S1 and S4). For each species page, this adjustment was made with a random set of pages in the Wikipedia language of that species page. For example, the Wikipedia page for *Panthera tigris* in the English language would be adjusted for a set of random pages in the English Wikipedia, whereas the page for *Panthera tigris* in French would be adjusted for a set of random pages in the French Wikipedia. To do so, we firstly calculated the rate of change for each random page in each language using ‘rlpi’, as in species pages. We then used a bootstrap resampling approach to calculate the average rate of change for all random pages in a given language at each timestep. The average rate of change in the random pages (*λ*_*rt*_) was calculated by bootstrapping the monthly rates of change 1000 times, and then extracting the bootstrapped mean. At each timestep, we then adjusted the species page rate of change by subtracting the monthly bootstrap estimated random rate of change (*λ*_*rt*_) as

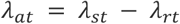

where *λ* = the rate of change, *t* = month, *r* = the bootstrapped trend for a given language, *s* = species page trend for that same language, and *a =* adjusted species trend.

For each species page the SAI is then where

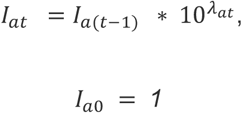

Where *I*_*at*_ = the species page SAI at time *t*.

To account for differences in the tortuosity of trends among Wikipedia languages (see Supplementary Information, Figure S7), we also smoothed the species page SAI in each Wikipedia language using a loess regression (span = 0.3), before transforming the smoothed species page SAI back into a rate of change.

After smoothing the species page SAI as above, we then calculated a species SAI for each species (across languages) by averaging rates of change at each time step across all languages. For example, the species *Panthera tigris* has the unique Wikidata ‘Q19939’, meaning the average rate of change in SAI for all species pages (irrespective of language) identified as ‘Q19939’ provides the overall rate of change for the species *Panthera tigris*.

We then calculated an overall SAI combining all species across 10 Wikipedia languages by averaging rates of change across all species SAIs. Bootstrap confidence intervals were calculated by taking the 2.5^th^ and 97.5^th^ percentiles of 1000 bootstrapped indices at each timestep. To check the extent to which single languages influence the overall SAI, we then jack-knifed the overall SAI for language, and removed any languages with a marked effect on the overall trend (see Supplementary Information, Figure S6).

Using the same approach as above, we also calculated an overall SAI for each taxonomic class for all languages combined, and each taxonomic class in each language. For each taxonomic class we again averaged the loess smoothed rate of change in species page SAI among languages, and then bootstrapped the species rate of change in SAI at each time step for each taxonomic class, as above. To check the extent to which single languages influence class level trends, we again jack-knifed the overall SAI for language, and removed any languages with a marked effect on the overall trend (see Supplementary Information, Figure S8). To calculate an overall SAI in each taxonomic class in each language, we bootstrapped the rate of change in species page SAI for the set of species pages in a given class-language combination.

### Predicting average monthly rate of change in the SAI

After calculating the SAI for all species pages on Wikipedia, we then calculated an average monthly rate of change in each smoothed species page SAI for the period January 2016 – January 2020. This average monthly rate of change was calculated across complete yearly periods to control for the effect of seasonality. To robustly explore whether change in awareness differs for various groups, we constructed 1 linear model and 2 mixed effects linear models predicting average monthly rate of change in species page SAI: 1) a linear model for average rate of change in species page SAI as a function of taxonomic class, language, and their interaction; 2) a mixed effects model for average rate of change in species page SAI as a function of taxonomic class, pollination contribution (Y/N), their interaction, and a random effect for language; and 3) a mixed effects models for average rate of change in species page SAI as a function of taxonomic class, traded status, and a random effect for language. Rather than attempting to find the most parsimonious model, we present full model predicted values, with AIC values for these and a set of candidate null models included in the Supplementary Information (Tables S8-S10).

## Results

### Wikipedia view dataset

Before removing incomplete series, our initial Wikipedia dataset included ∼2.23 billion page views for IUCN species across the Arabic, Chinese, English, French, German, Italian, Japanese, Portuguese, Russian, and Spanish Wikipedias. These views were represented over a period of 1735 days between the 1st July 2015 and 31st March 2020. Views for each language varied from ∼24.92 million views in the Arabic Wikipedia to ∼1.08 billion views in the English Wikipedia (Figure 2). For all languages unique species number was highest in the ray-finned fishes at 13,571 and lowest in the insects at 2,743 (Figure 2, see Supplementary Information Figure S1 for a full language breakdown). After subsetting for series represented for every month, the proportion of complete series was lowest in the Arabic Wikipedia, specifically the ray-finned fishes (∼35%) and the reptiles (∼38%). Most taxonomic classes for most languages had complete series in at least 80% of the species in that grouping (Supplementary Information, Figure S1).

**Figure 2.**
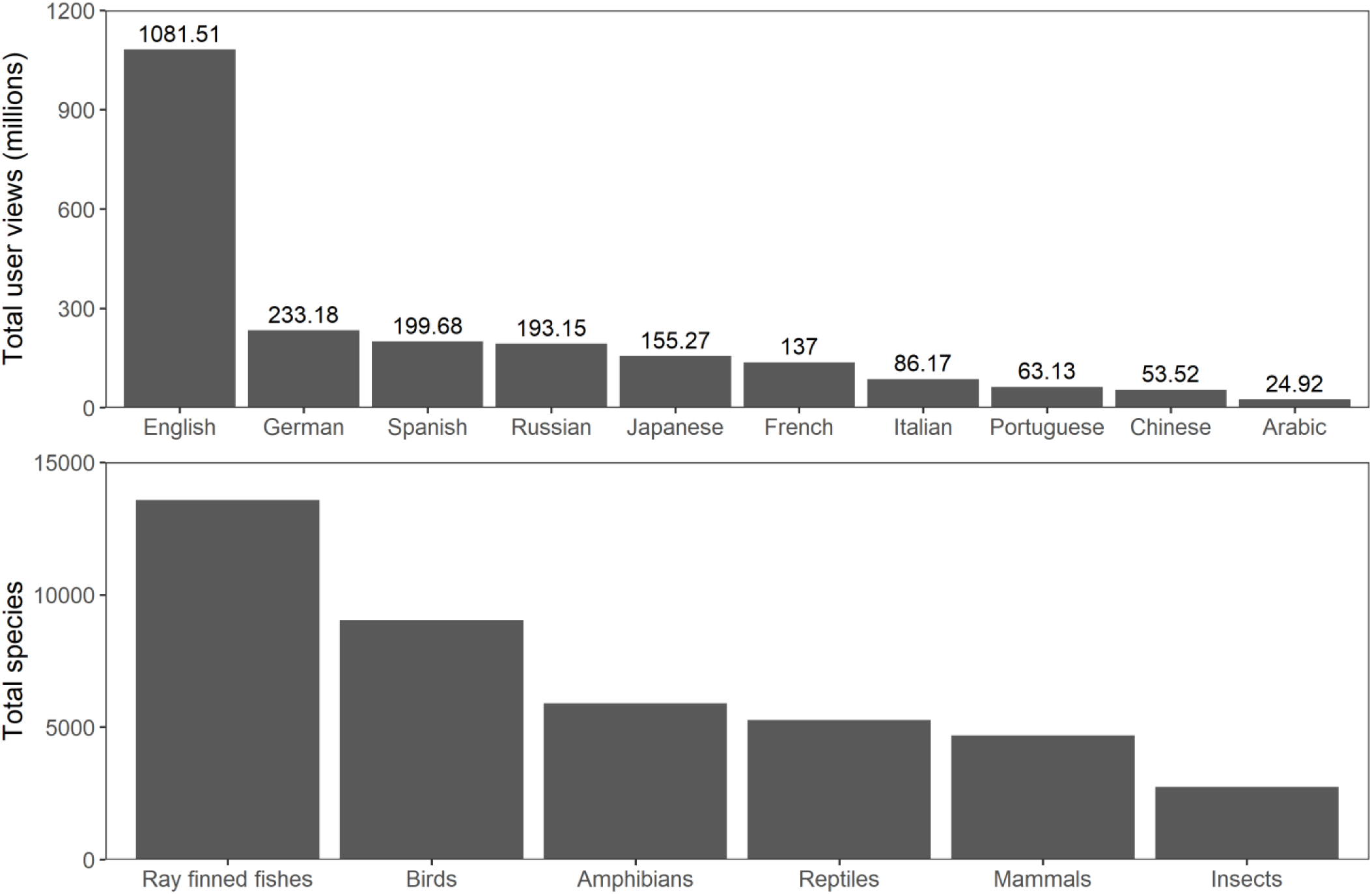
The total number of views and unique species for the initial Wikipedia view dataset, before removing incomplete series. Top: the total number of views for IUCN species in each Wikipedia language, in order of magnitude: English, German, Spanish, Russian, Japanese, French, Italian, Portuguese, Chinese, and Arabic. Bottom: the total number of unique IUCN species with Wikipedia pages in each of 6 taxonomic classes (ray-finned fishes, birds, amphibians, reptiles, mammals, and insects), across all 10 languages of the top panel.

After removing pages also present in the species set, our set of random views consisted of ∼2.82 billion views across 113,622 random pages (Supplementary Information, Figure S3), again for the same 1735 day period. The total number of random views was highest for the English Wikipedia at ∼629.85 million views, and lowest in the Arabic Wikipedia at ∼87.94 million views (Supplementary Information, Figure S3). After subsetting for only random pages represented for all months, total random pages varied from 3486 in the Arabic Wikipedia to 9174 in the Japanese Wikipedia (Supplementary Information, Figure S4).

### Absolute awareness of biodiversity

Among taxonomic classes, reptiles have consistently higher absolute awareness, appearing in the top 2 classes for 7/10 languages (Supplementary Information, Figure S11). Amphibians on the other hand have consistently lower awareness, appearing in the bottom 2 classes for 8/10 languages. Some languages appear to have uniquely high absolute awareness for specific classes. For example, the ray-finned fish have the highest absolute awareness in the Japanese Wikipedia (Supplementary Information, Figure S11). Across all languages, absolute awareness (total views) is significantly higher in traded species (Figure 2; F = 15206.44, p < 0.001, Table S2), but not significantly different in pollinating species (F = 0.3869, p = 0.5339, Table S1).

### Awareness Index (SAI)

The overall SAI for all taxa and languages was markedly affected by the inclusion of the French Wikipedia (see Supplementary Information Figure S6), meaning it was excluded from analyses presenting aggregated change at the overall level. With the exclusion of the French Wikipedia, the overall SAI increased from July 2015-March 2020, with marked declines in mid 2016 and mid 2018 (Figure 4).

**Figure 3.**
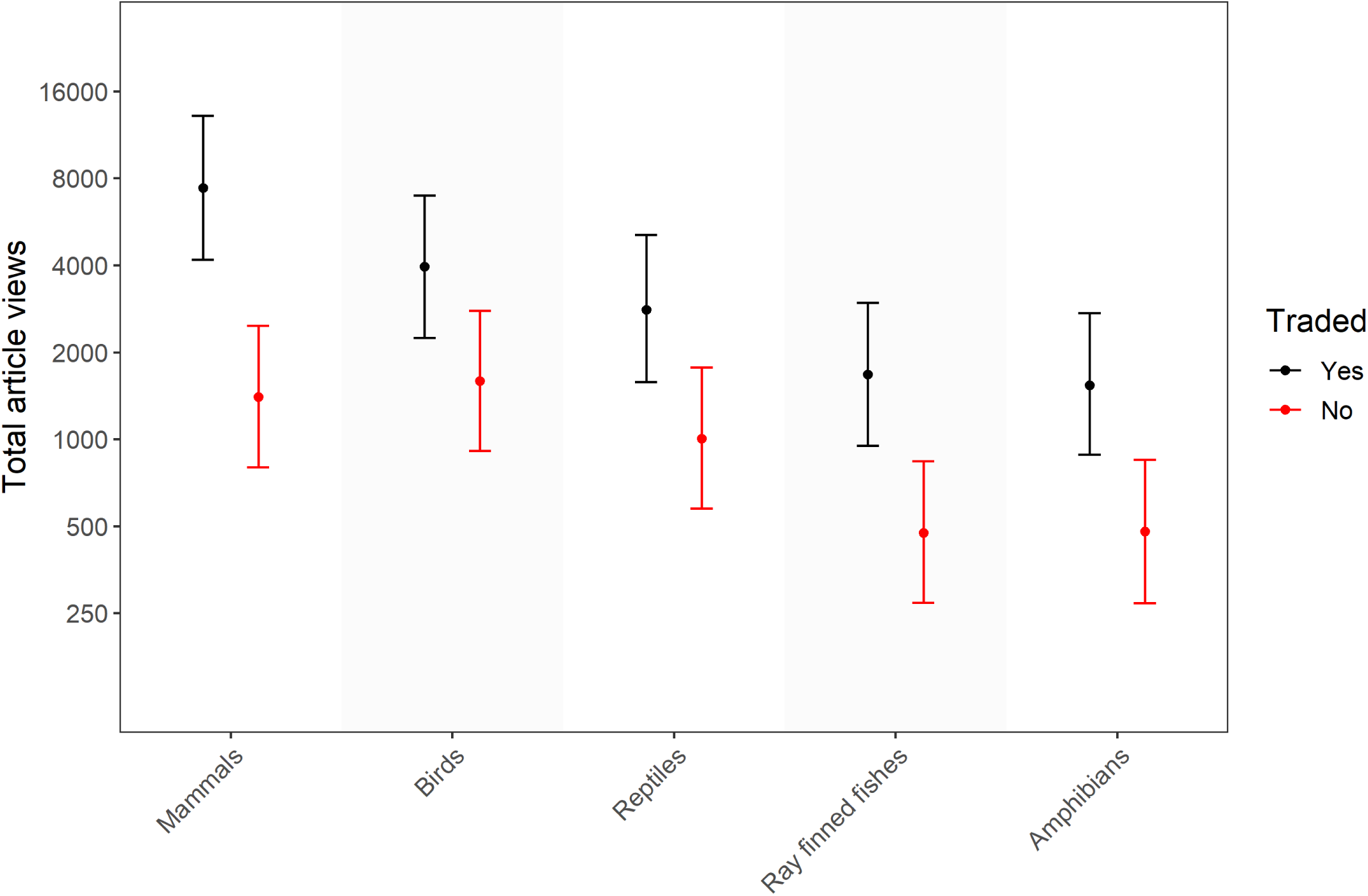
Total Wikipedia article views for traded and non-traded species across 6 taxonomic classes (mammals, birds, reptiles, ray-finned fishes, and amphibians) and 10 Wikipedia languages (Arabic, Chinese, English, French, German, Italian, Japanese, Portuguese, Russian, Spanish). Predicted values were generated using a generalised linear mixed-effects model, modelling total article views as a function of the fixed effects taxonomic class, the presence of trade (Y/N), and their interaction, and the random effect language. Effect sizes were calculated by drawing fixed effects 1,000 times based on the variance-covariance matrix, and then calculating the median value (shown as points), and 2.5^th^ and 97.5^th^ percentiles (shown as error bars). Black error bars represent species that *are* known to be traded, and red error bars represent species that *are not* known to be traded or harvested. Taxonomic classes are ordered by the magnitude of total article views, from highest on the left to lowest on the right.

**Figure 4.**
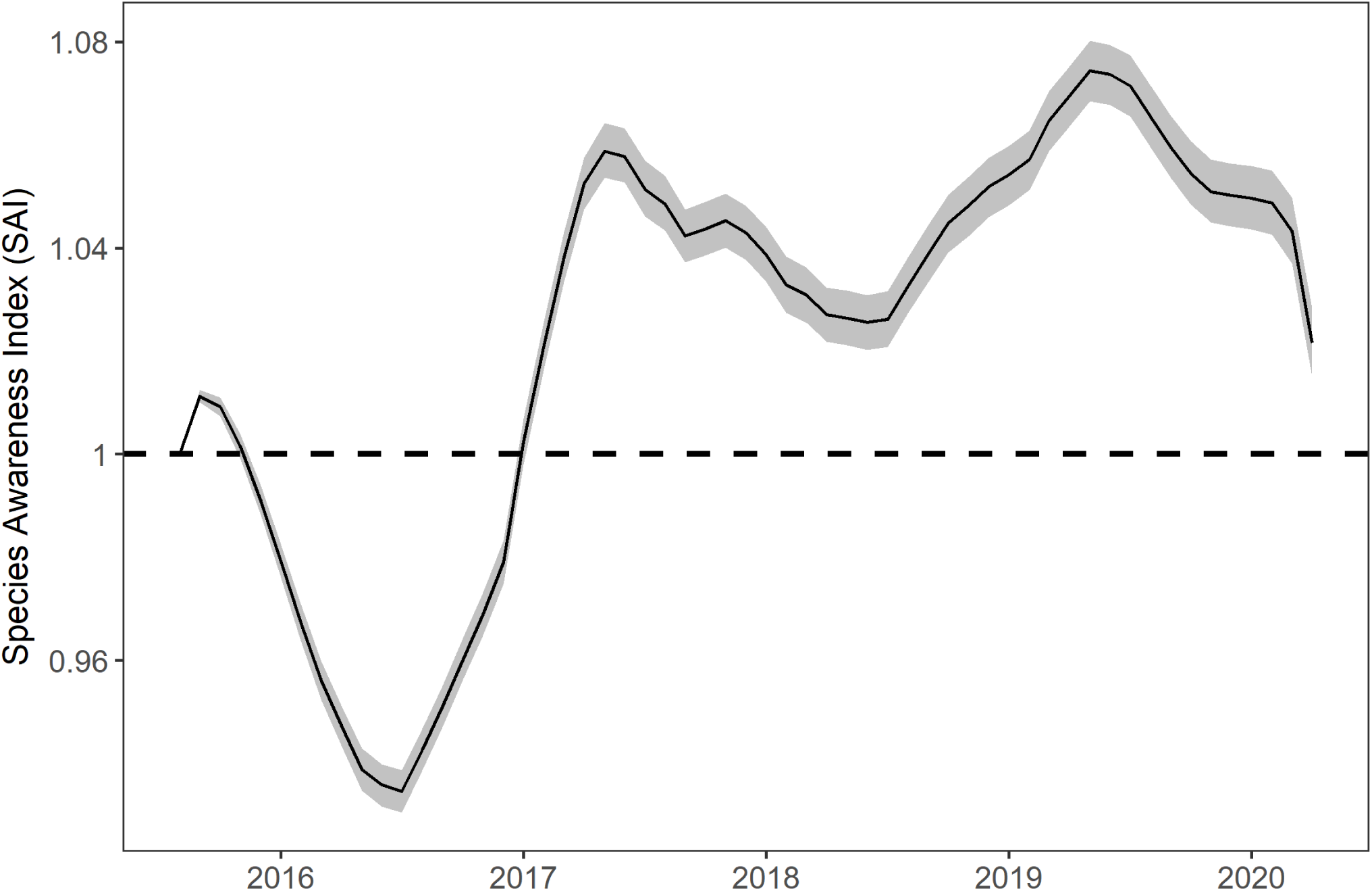
The Species Awareness Index (SAI) for 6 taxonomic classes (reptiles, ray-finned fish, mammals, birds, insects, and amphibians) and 9 Wikipedia languages (Arabic, Chinese, English, German, Italian, Japanese, Portuguese, Russian, and Spanish) for the period July 2015-March 2020. The French Wikipedia was removed here given its marked influence on the aggregated SAI (see Supplementary Information, Figure S6). Here the black line represents the mean of the bootstrapped indices at each monthly timestep, and the grey band the 2.5^th^ and 97.5^th^ percentiles.

At the level of taxonomic class, jack-knifing trends by language again showed that the French Wikipedia was markedly affecting the overall trend (see Supplementary Information Figure S8). With the exclusion of the French Wikipedia, over the period July 2015-March 2020 all of the reptiles, ray-finned fish, mammals, and birds appear to have increased in awareness, whilst the amphibians and insects appear to have decreased (Figure 5). Birds experienced a peak in early 2017 (Figure 5), driven by an increase across multiple languages (Figure 6). The mammals appear to be experiencing a consistent and steady increase in awareness, particularly in the Japanese Wikipedia (Figure 6). The amphibians and insects both experienced a pronounced drop in awareness from the start of the series to mid 2016 before increasing, the cause of which is unclear. The trend for both the reptiles and insects is highly seasonal for multiple languages, peaking in July-August of each year, with the notable exception of the English language for insects (Figure 6).

**Figure 5.**
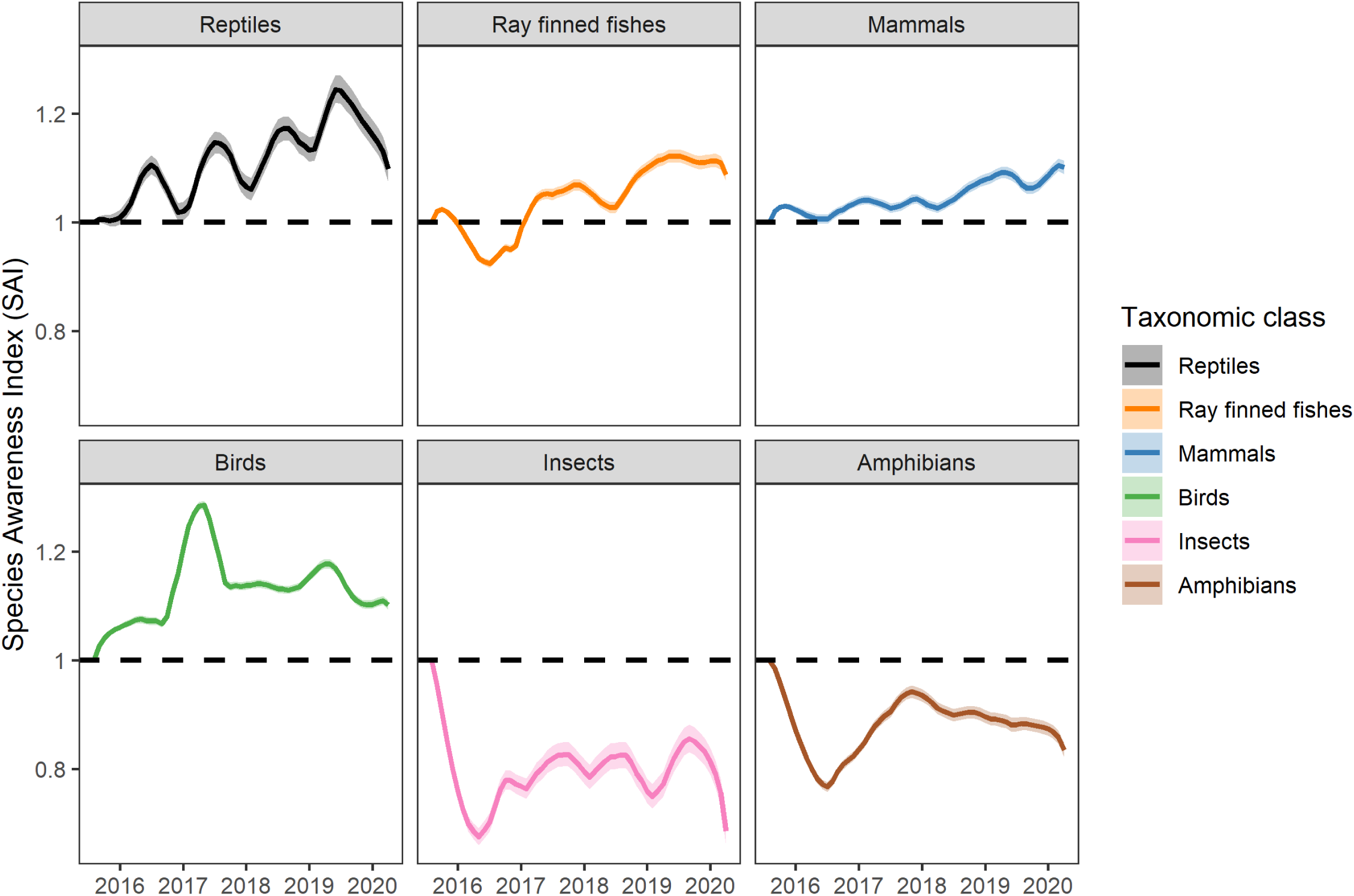
The Species Awareness Index (SAI) for 6 taxonomic classes (reptiles, ray-finned fishes, mammals, birds, insects, and amphibians) and 9 Wikipedia languages (Arabic, Chinese, English, German, Italian, Japanese, Portuguese, Russian, and Spanish), for the period July 2015-March 2020. The French Wikipedia was removed here given its marked influence on the aggregated SAI (see Supplementary Information, Figure S8). Coloured lines represent the mean of the bootstrapped indices at each monthly time step, and coloured bands the 2.5^th^ and 97.5^th^ percentiles: reptiles (black), ray-finned fishes (orange), mammals (blue), birds (green), insects (pink), and amphibians (brown). Taxonomic class panels are ordered by the magnitude of overall increase in each taxonomic class.

**Figure 6.**
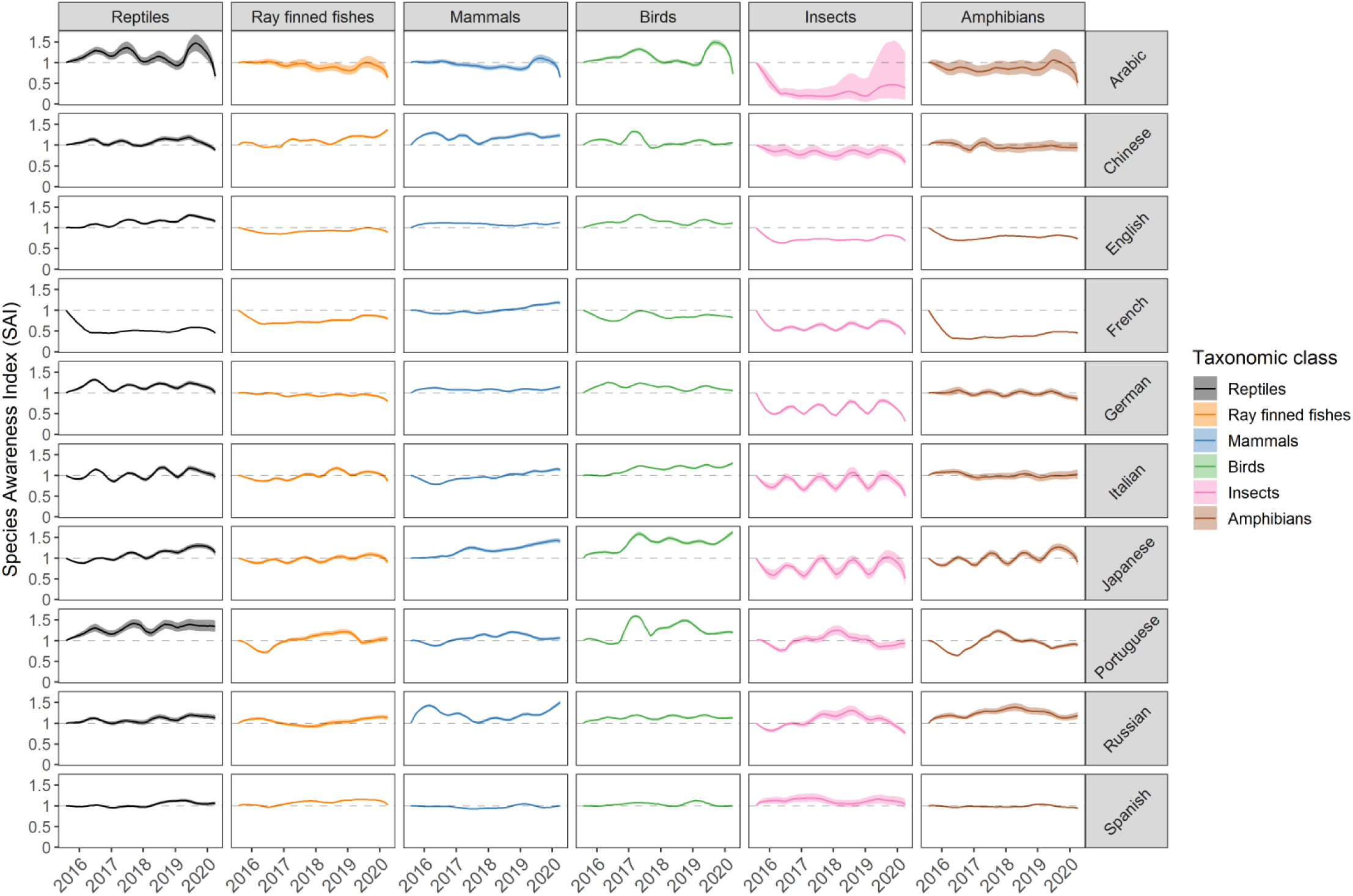
The Species Awareness Index (SAI) for 6 taxonomic classes (reptiles, ray-finned fishes, mammals, birds, insects, and amphibians) across 10 Wikipedia languages, for the period July 2015-March 2020. Coloured lines represent the mean of the bootstrapped indices at each monthly each time step, and coloured bands the 2.5^th^ and 97.5^th^ percentiles: reptiles (black), ray-finned fishes (orange), mammals (blue), birds (green), insects (pink), and amphibians (brown). Taxonomic class panels are ordered by the magnitude of overall increase in each taxonomic class, and for language alphabetically.

### Predicting average monthly rate of change in species page SAI

Average monthly rate of change in species page SAI for the period January 2016-January 2020 differed significantly for all of taxonomic class, language, and their interaction (Figure 7, Table S5). At the level of taxonomic class, the reptiles and ray-finned fishes are increasing in awareness the fastest, and the insects and amphibians are either increasing slowly or declining (with the exception of the Japanese Wikipedia). Among languages, rate of change in species page SAI is highest in the Japanese and Portuguese Wikipedias, and lowest in the German and Chinese Wikipedias. Although absolute interest is significantly greater in traded species (See Figure 3, Table S2), over the period January 2016-January 2020 average monthly rate of change in the species page SAI appears not to be related to either trade status (Figure S14, Table S4) or pollination contribution (Figure S14, Table S3).

**Figure 7.**
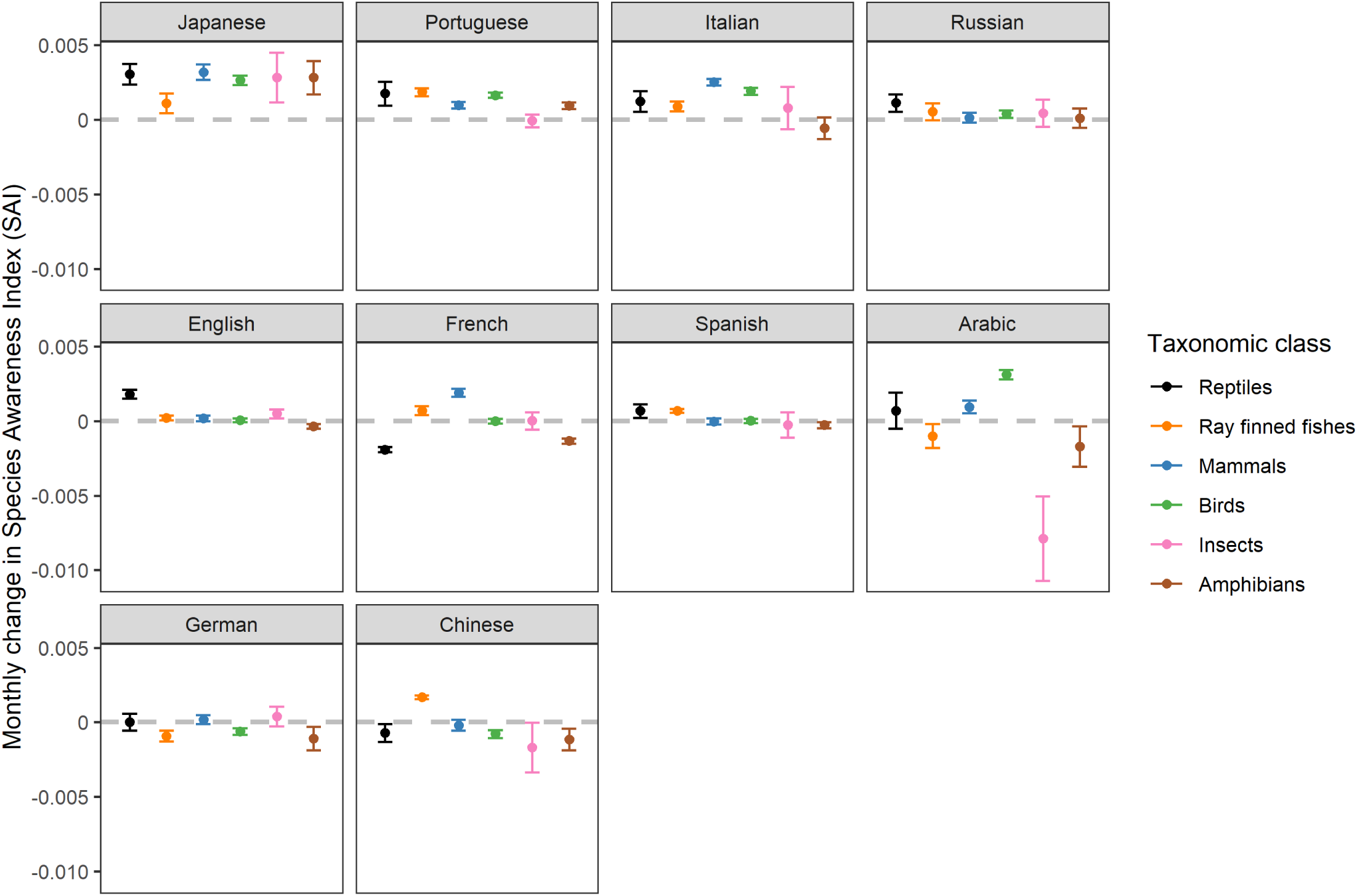
Average monthly rate of change for the species page SAI for 6 taxonomic classes across 10 Wikipedia languages. Errors bars represent the predicted values of a linear model, fitting average monthly change in the species page SAI as a function of taxonomic class, Wikipedia language, and their interaction. Fitted values were generated from the linear model with the R function predict (represented as points), and 95% confidences intervals from the fitted values +/-1.96 * standard error. The colour of error bars refers to taxonomic class: reptiles (black), ray-finned fishes (orange), mammals (blue), birds (green), insects (pink), and amphibians (brown).

## Discussion

In this study we introduced the Species Awareness Index (SAI), an index of change in public awareness of biodiversity. The SAI is derived from views of individual species pages, enabling disaggregation by a variety of variables, such as taxonomy, language, geographic distribution, and ecosystem service provision. We show that awareness of biodiversity overall has increased, with increases highest in the reptiles and ray-finned fish. Although biodiversity awareness is increasing overall, we also show that some groups (i.e. the amphibians and insects) are decreasing or only marginally increasing in awareness. Among languages, increases are highest in the Japanese and Portuguese Wikipedias, and lowest in the Chinese and German Wikipedias. Although our results suggest that awareness of biodiversity has increased since July 2015, at this timescale increases do not appear to be related to the trade of species or animal pollination. As an indicator for biodiversity awareness, a Wikipedia derived metric such as the SAI represents a useful additional data source, given its explicit and unambiguous link to biodiversity itself at multiple scales.

The link between culture and perceived biodiversity value or awareness is widely recognised (Daniel et al 2012; Cooper et al 2019; Roll et al 2016; Ladle et al 2019), but for a culturomics metric such as the SAI the drivers and implications of change are complex. Overall trends capture many different drivers of awareness, making it difficult to isolate the causes for a given increase or decrease. The Chinese Wikipedia, for example, shows a consistent decrease in awareness for 5 taxonomic classes, but a consistent increase for ray-finned fish. We hypothesised that this increase for ray-finned fish may be driven by increasing fish consumption, since seafood demand in China has increased significantly in recent years (FAO, 2020). However, in a brief additional analysis, we found no significant difference between the rate of change for traded and non-traded ray-finned fish in the Chinese Wikipedia (Supplementary Information, Figure S13). This would indicate that the greater rate of change for the Chinese ray-finned fish may not be driven by consumption alone. The Japanese Wikipedia is also of note, given its consistent increase in awareness across all 6 taxonomic classes. This awareness increase is concordant with the results of conventional surveys, in which Japan has amongst the largest percentage point increases for familiarity with the term ‘biodiversity’ (UEBT 2019). However, it is unclear what may be driving this change in awareness. Counterfactual scenario modelling could help to better understand such relationships, previously demonstrated in a number of recent conservation culturomics studies (Acerbi et al 2020; Verissimo et al 2020; Fernandez-Bellon & Kane 2019).

Change in biodiversity awareness reflected by the SAI is related to, but largely distinct from absolute awareness. For example, although awareness of reptiles may be rising fastest, this does not necessarily mean that absolute awareness is also high. Rather, one might expect that groups of high absolute awareness would often be stable or increasing slower, since the margin within which they can continue to increase is smaller. In the Japanese Wikipedia, for example, ray-finned fish have the highest absolute awareness, but the lowest average monthly rate of change in species page SAI. Future work should further explore the relationship between absolute and change in awareness, as well as the influence of other ecosystem services such as pest control and predation on awareness. It would also be interesting to include variables previously applied in absolute awareness research, such as threat status and venomosity (Roll et al 2016), and geographic range and phenotypic distinctiveness (Ladle et al 2019).

Although some taxa may have experienced an increase in interest on Wikipedia, this does not necessarily mean a sustained increase in awareness or knowledge of biodiversity. Our index can show whether views increased or decreased on aggregate for a given grouping, from which we can infer exposure to species-related information. However, we cannot tell why a given page was visited, or whether information related to that page was retained. Previous research has shown how on-site Google Analytics can be used to return a suite of metrics intimating as to the reasons for a given visit (Soriano-Redondo et al 2016), such as the site used to reach a page, but for Wikipedia this data is not publicly available. Text-mining could help to quantify the type of information users are exposed to on Wikipedia. For example, calculating a text similarity for each species page to a reference text on pollination would provide an indication as to the pollination salience of a given species page. Similar approaches have been applied in the context of climate change and invasive species, using rate of threat-related terms as an indicator of threat salience (Jaric et al 2020). Given opinions on species within a taxonomic class can be so diverse (Sumner et al 2018), such an approach could help to distinguish between drivers of awareness.

There are other limitations associated with the SAI. First, the short length of our Wikipedia time series presents a problem for the interpretation of our index. Future iterations of the SAI will aim to include views archived from before 2015, which presents additional problems in reconciling views aggregated across multiple formats. Second, an index based on internet activity will only be representative of those that have access to the internet, and use the internet to access Wikipedia. Given ∼50% of the global population has access to the internet (The World Bank, 2020), and ∼15% of those regularly access Wikipedia (Graham et al., 2014), the SAI cannot be globally representative. In mainland China for example, the dominant online encyclopedia is Baidu Baike, with Wikipedia views for the Chinese (zh) Wikipedia coming predominantly from Taiwan and Hong Kong (Wikimedia Traffic Analysis Report, 2018). Future iterations of the SAI could incorporate page views for species on Baidu Baike, although this presents problems in requiring a distinct computational approach.

Although the SAI and Biodiversity Engagement have value as independent metrics, as in Cooper et al. (2019) we emphasise the importance of multiple online platforms for inferring public biodiversity awareness. Particularly since the SAI provides an explicit link to biodiversity itself, its inclusion could provide a more holistic understanding as to how biodiversity awareness is changing. Combining the SAI with the Biodiversity Engagement Indicator presents two core challenges. First, both the Biodiversity Engagement Indicator and SAI are measured on different units, inherent to their underlying methodology. The Biodiversity Engagement Indicator is scaled on a 0-100 scale in a manner analogous to Google Trends. The SAI on the other hand is scaled relative to a benchmark index of 0, in an approach inspired by the LPI. Second, there are problems of geographic scale in combining the Biodiversity Engagement Indicator and SAI. Namely, the Biodiversity Engagement Indicator is aggregated at the country-level, whereas the SAI cut across countries at the language-level.

Given differences in units and geographic scale, combining a Wikipedia metric and the Biodiversity Engagement Indicator is not simple. One potential solution could be to rescale the SAI on a 0-100 scale, disaggregate by language, and then calculate a weighted average among languages to reflect the proportion of users for a given country (Figure 8). The data for calculating such a weighting language level trend is provided by Wikipedia, in a format amenable to web-scraping (Wikimedia Traffic Analysis Report, 2018). Such an approach would solve both the problem of differing units and geographic scale, transforming the SAI into a national metric amenable to averaging alongside a Twitter, Newspaper, and Google Trends score (Figure 8).

**Figure 8.**
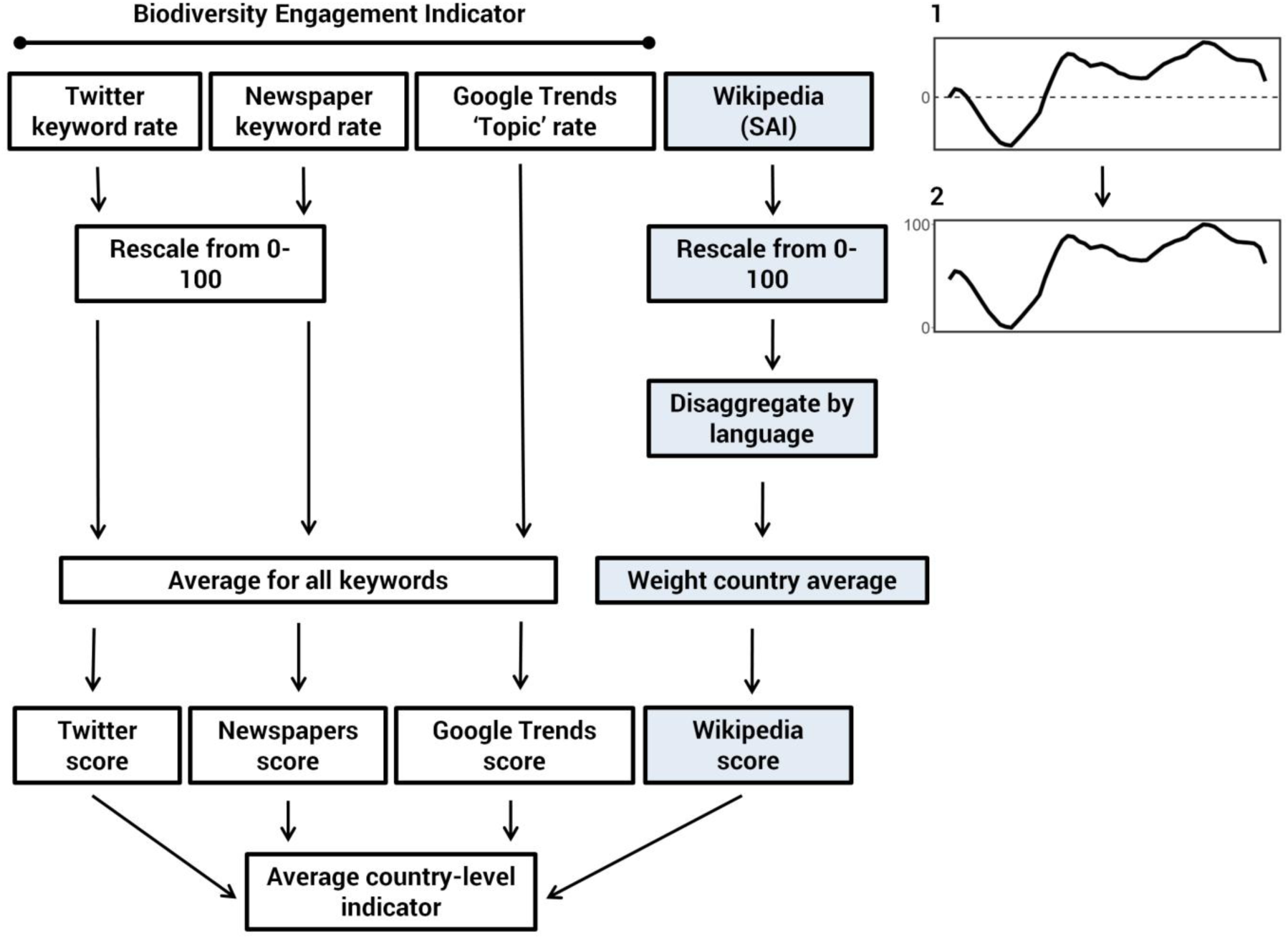
Schematic adapted from Cooper et al (2019), demonstrating one method through which the Species Awareness Index (SAI) might be incorporated into the Biodiversity Engagement Indicator for a single indicator of country-level biodiversity awareness. All text squares in white represent the methodological pathway for the original Biodiversity Engagement Indicator, and all squares in pale blue represent a potential additional pathway for combining the SAI. Box 1 (top right) represents the overall SAI scaled starting at 0. Box 2 represents the overall SAI rescaled between 0-100, consistent with the Biodiversity Engagement Indicator.

As global internet penetration increases and biodiversity continues to experience decline, digital metrics for public biodiversity awareness will become both more informative and more important. Here we presented the Species Awareness Index (SAI), a metric derived from Wikipedia views depicting change in awareness for biodiversity online. We used this metric to show that overall awareness of biodiversity is increasing marginally, although this increase is inconsistent among taxonomic groups and languages. We also showed that such increases appear not to be related to either the pollination contribution or trade status of a species. We concluded by suggesting one approach through which the SAI could be combined with the Biodiversity Engagement Indicator, providing a more holistic understanding of public biodiversity awareness in the digital realm.

## Footnotes

1 https://wikitech.wikimedia.org/wiki/Analytics/AQS/Pageviews

2 https://www.mediawiki.org/wiki/API:Main_page (using ‘wbgetentities’)

3 https://www.mediawiki.org/wiki/API:Random

