## Supporting information for "The Species Awareness Index (SAI): a Wikipedia-derived conservation culturomics metric for public biodiversity awareness"

**Extended methodology for pollinator dataset construction**

We first created a list of possible pollinating animal genera through automatic text analysis of the pollination literature. We considered the pollination literature to be any primary research article published in English returned through a search for the term “pollinat*” in Scopus, and which mentioned an animal species in the abstract. We considered a possible pollinating genus to be any animal genus appearing as part of a Latin binomial in a pollination-related abstract returned from Scopus. Genera scraping was accomplished using the *Taxonfinder* and *Neti* *Neti* algorithms implemented in the ‘taxize’ R package (CRAN 2018), with animal species confirmed through a series of character string matches to the COL (see Millard et al 2020 for a detailed methodology).

For each possible pollinating genus, we then read the abstracts in which these animals appeared, searching for evidence confirming that genus as pollinating. For any situation in which the abstract was inconclusive, we also searched the full text of the paper for more definitive evidence. For each confirmed pollinating genus, we then assigned a level of experimental confidence between 1 and 4 based on the type of evidence (following (Ollerton and Liede 1997)): 1) experimental evidence confirming pollination; 2) evidence of pollen carrying; 3) evidence of nectar/pollen feeding; 4) evidence of non-destructive/non-predatory flower visitation. Our process of assigning evidence was one of maximisation: we read abstracts for each genera searching for the highest level of evidence, either until we could be sure that the confidence value should be 1, or we ran out of abstracts for that genera. Non-destructive flower visitor refers to any animal which visits a flower without causing damage to the plant. This meant the exclusion of most ants, which are typically referred to as poor pollen vectors, given that they damage pollen through secretions from the meta-pleural gland (Dutton and Frederickson 2012). Non-predatory flower visitor refers to any animal which visits for some purpose other than predation. This meant the exclusion of animals such as crab spiders, which predate on pollinators during visitation, and therefore contribute minimally or negatively to pollination (Dukas and Morse 2003). We did not classify broad statements as evidence for pollination—for example, one study stated that *Phylidonyris novaehollandiae* is a “key pollinator” (Myers et al 2012)—unless it was associated with specific evidence reinforcing that statement, or some claim that pollination in that genus is “well-known” or “widely acknowledged”.

Given that we only had direct evidence for a sample of all pollinating genera, we then searched for higher-level clades of pollinators. From the confirmed pollinators in the original list of genera, we identified all unique families with at least one pollinator. For each family, we assessed the breadth of evidence for pollination through consulting the abstracts, taxonomic clade reference books, and expert opinion. For any family with evidence of pollination across multiple branches of that family, and no evidence of any species definitely not pollinating, we assumed that the whole family is pollinating. If unable to extrapolate across the whole family, we then searched progressively lower taxonomic clades (i.e. subfamily, tribe, subtribe), searching for the point at which we could be relatively confident that the entire clade contributes to pollination. If unable to extrapolate for a given clade, we kept only the genera with direct evidence. For example, within the family Formicidae (ants), we found no clade across which we could confidently extrapolate, and so kept only those genera for which we had direct evidence. We assigned a level of confidence for these extrapolated clade designations, which varied between 5.1 and 5.4, reflecting the quality of evidence available for most species in that clade, with the ‘5’ indicating an extrapolated clade: 5.1, experimental evidence across multiple groups within that clade; 5.2, evidence of pollen carrying across multiple groups; 5.3, evidence of nectar/pollen feeding across multiple groups; 5.4, evidence of non-destructive/non-predatory flower visitation across multiple groups. After checking all families represented by the original list of genera, we then inspected additional resources, searching for any pollinating families we may have missed. Each of these resources included evidence of only nectar/pollen feeding or flower visitation, meaning all families extrapolated through additional resources were assigned a confidence level of either 5.3 or 5.4.

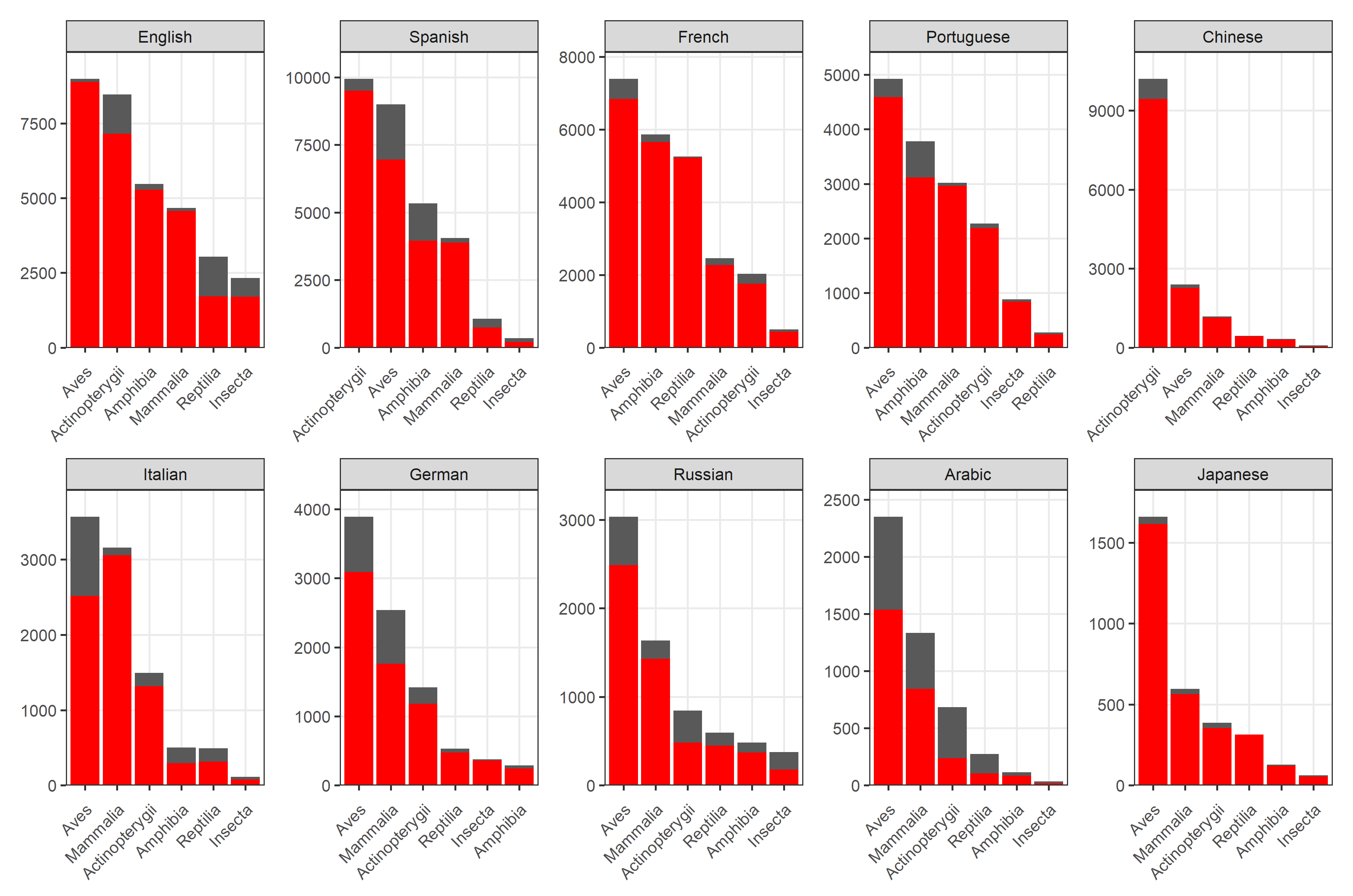

Figure S1. The number of complete Wikipedia view series for each taxonomic class in each language. Red bars represent species with complete series (i.e. data for each month over the period July 2015-March 2020), from which the SAI was calculated. Black bars represent the total species for each taxonomic class in each language, including species for which the series of views is incomplete.

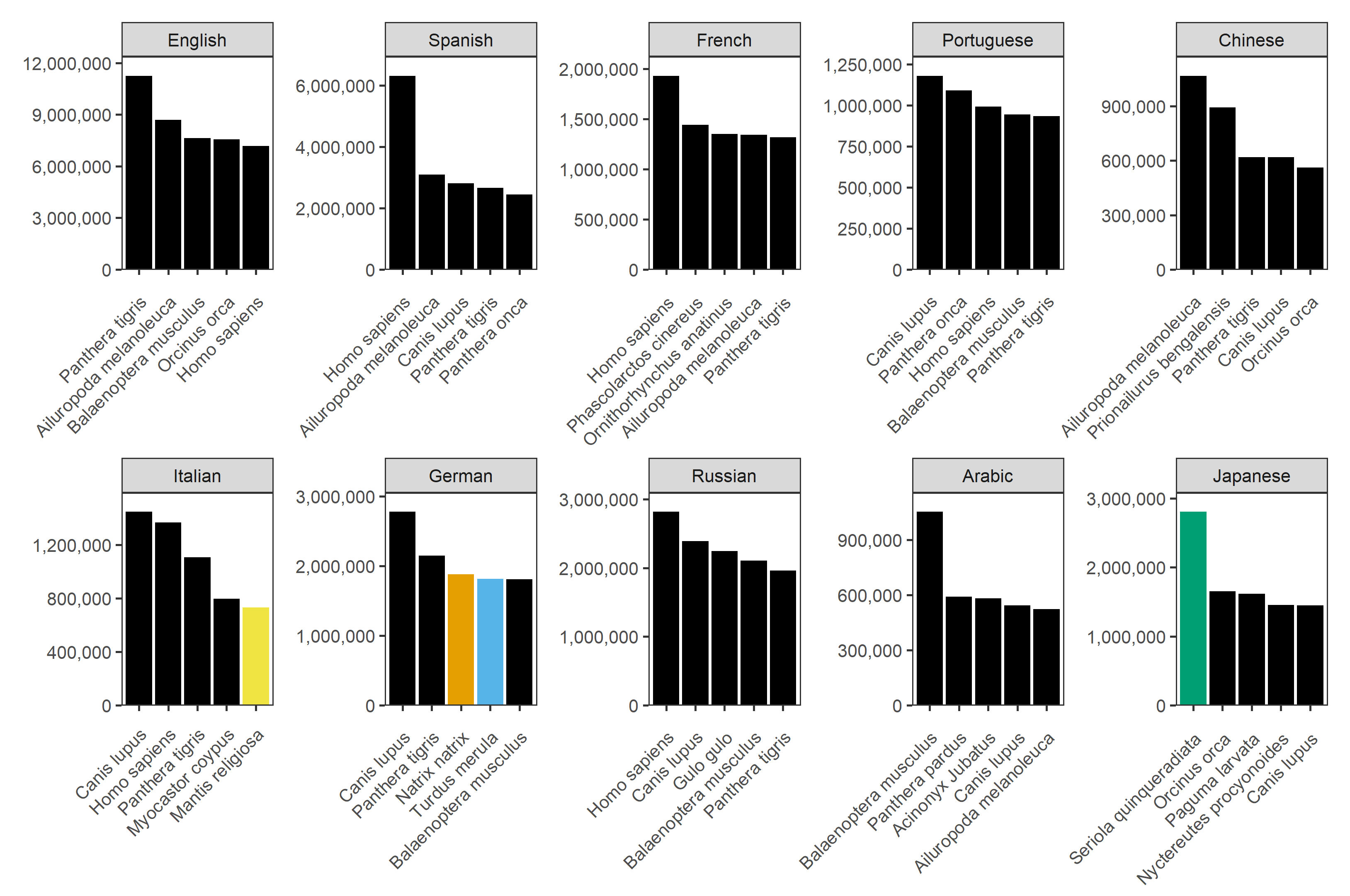

Figure S2. The top 5 viewed species in each Wikipedia language. Each colour fill refers to a taxonomic class, consistent across all 10 panels: black (mammals), yellow (insects), green (ray-finned fish), orange (reptiles), and blue (birds).

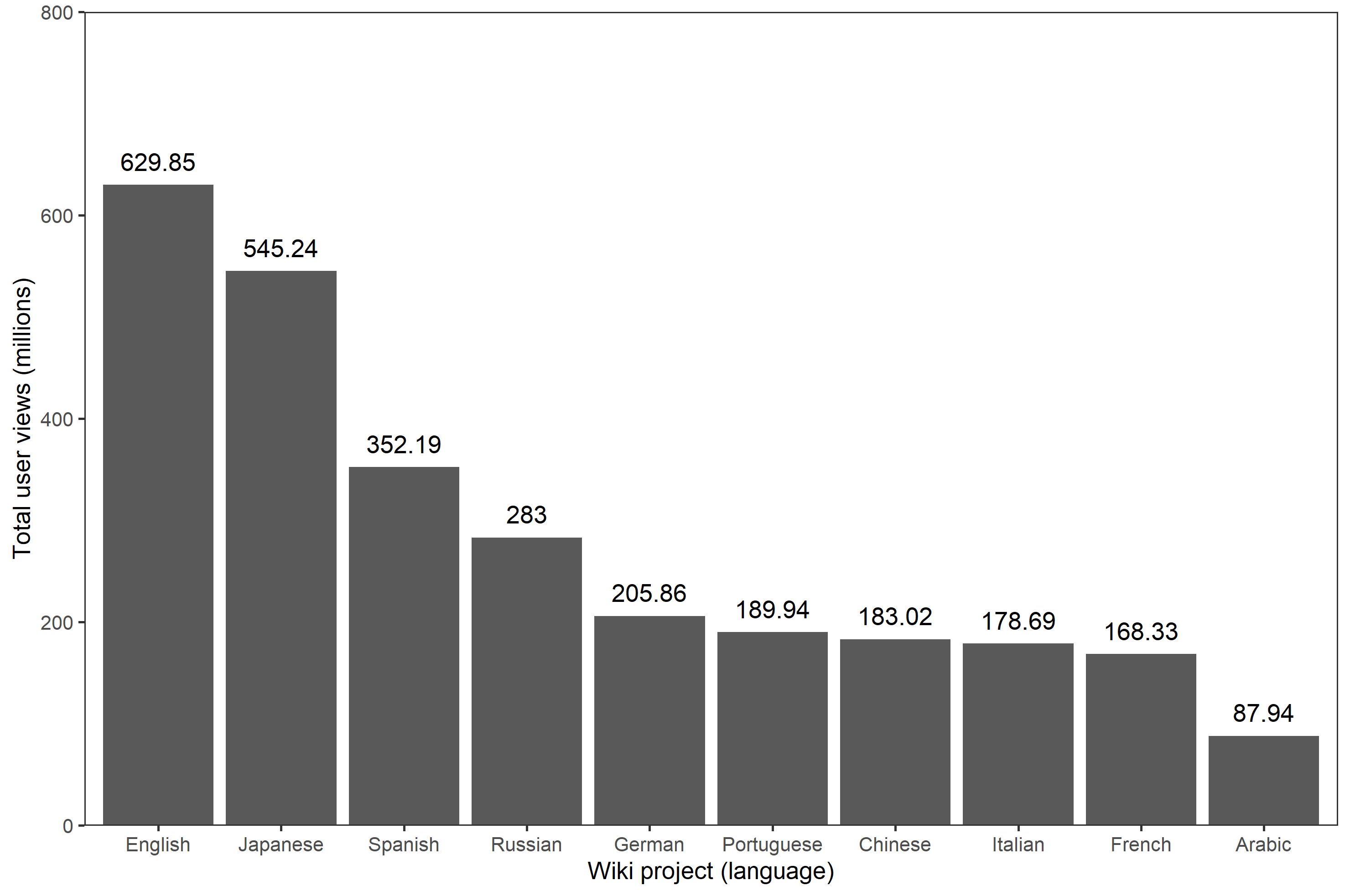

Figure S3. The total number of Wikipedia views for the set of random pages in each language, before removing incomplete series.

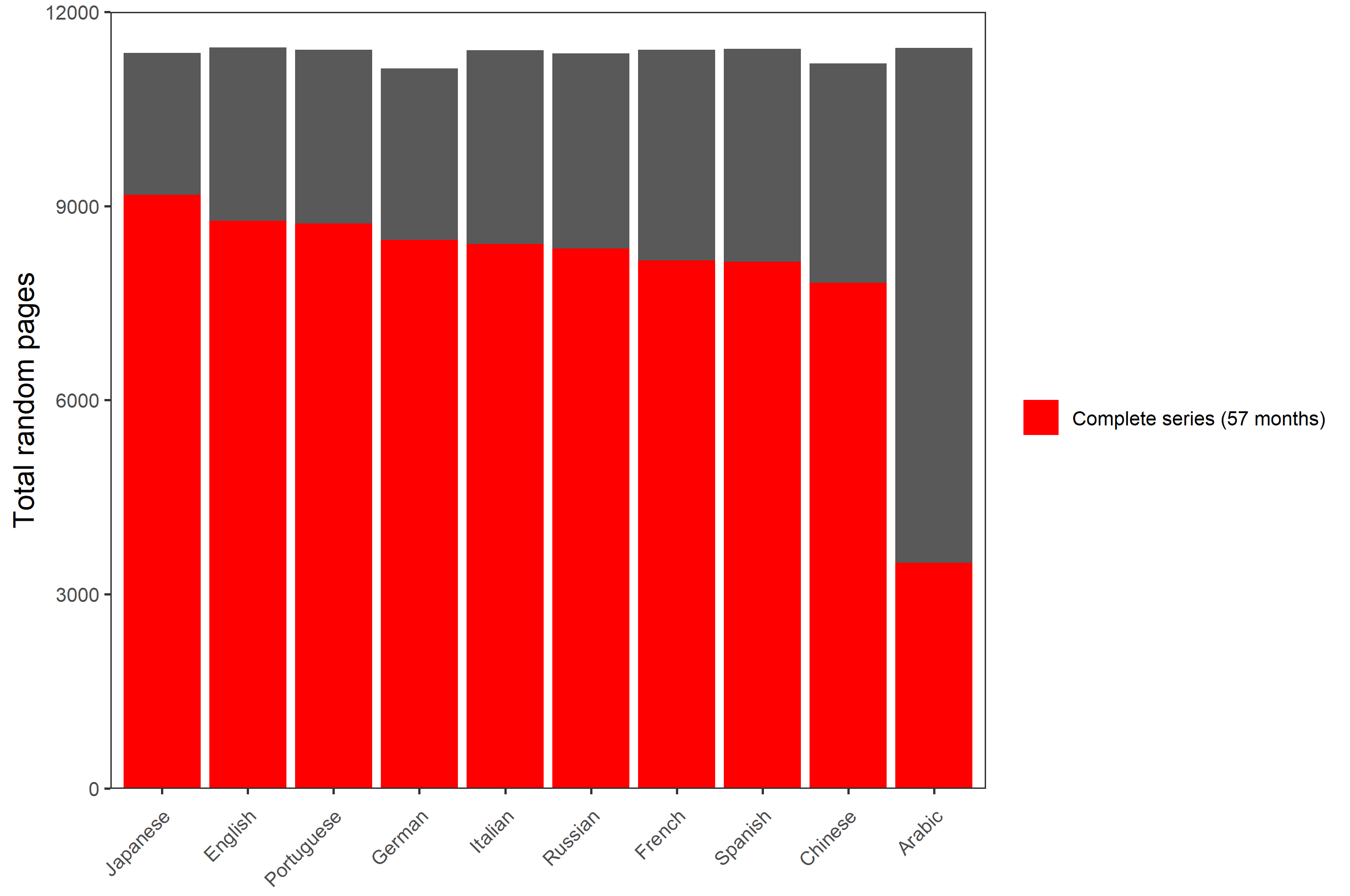

Figure S4. The total number of complete series pages for each set of random pages in each language. Red bars represent pages with complete series (i.e. data for each month over the period July 2015-March 2020), from which the random index was calculated. Black bars represent the total number of random pages in each language, including pages for which the series of views is incomplete.

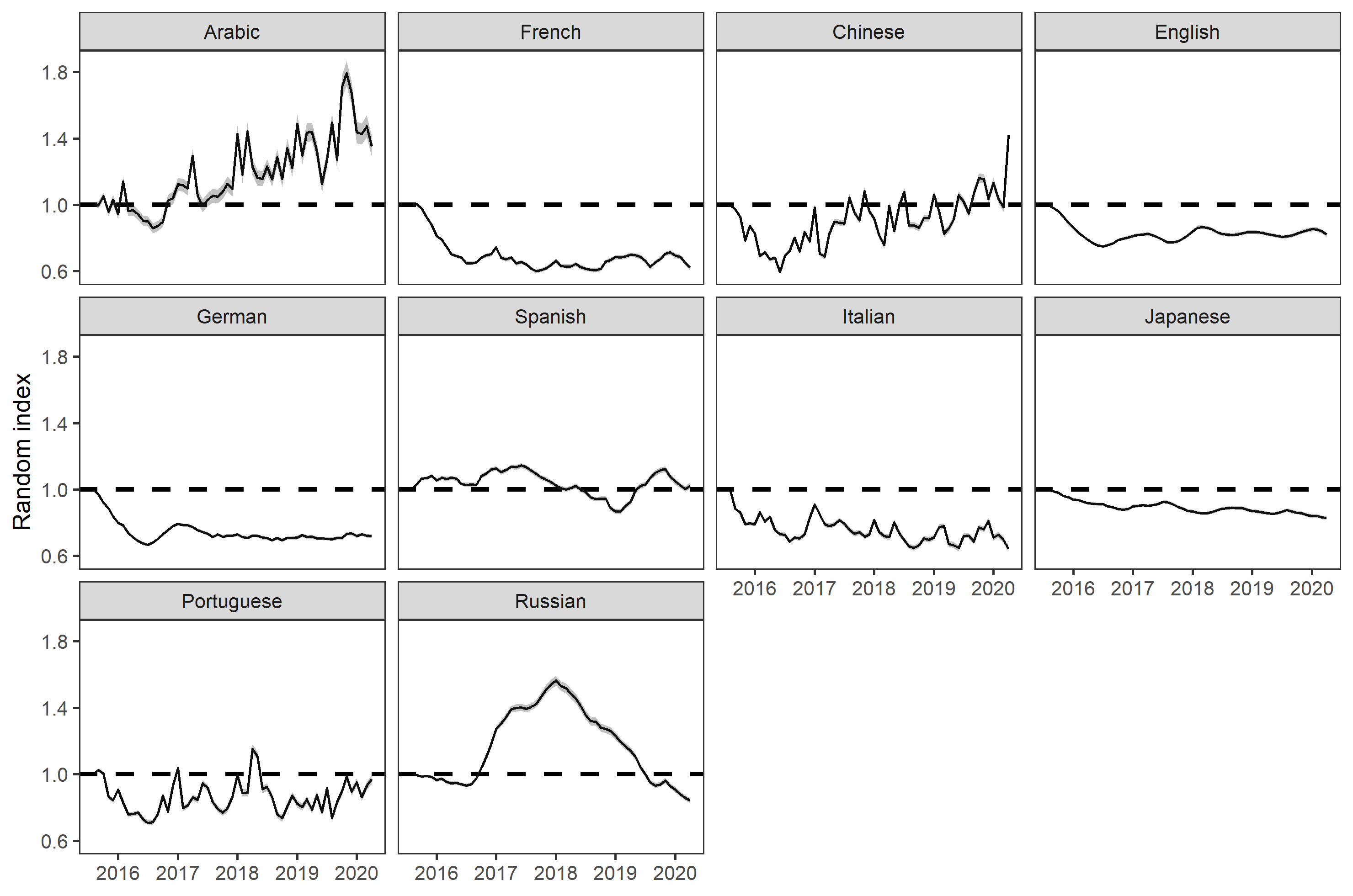

Figure S5. The random index for the set of complete series random pages in each language. Black lines represent the mean of the bootstrapped indices at each timestep, and grey bars the 2.5^th^ and 97.5^th^ percentiles. The random index provides an estimate of the overall change in popularity for each Wikipedia language (i.e. a decrease in the random index for a given language would imply that the popularity of Wikipedia in that language has decreased). To calculate the species page SAI, the rate of change in the mean random index is subtracted from the rate of change in each species page.

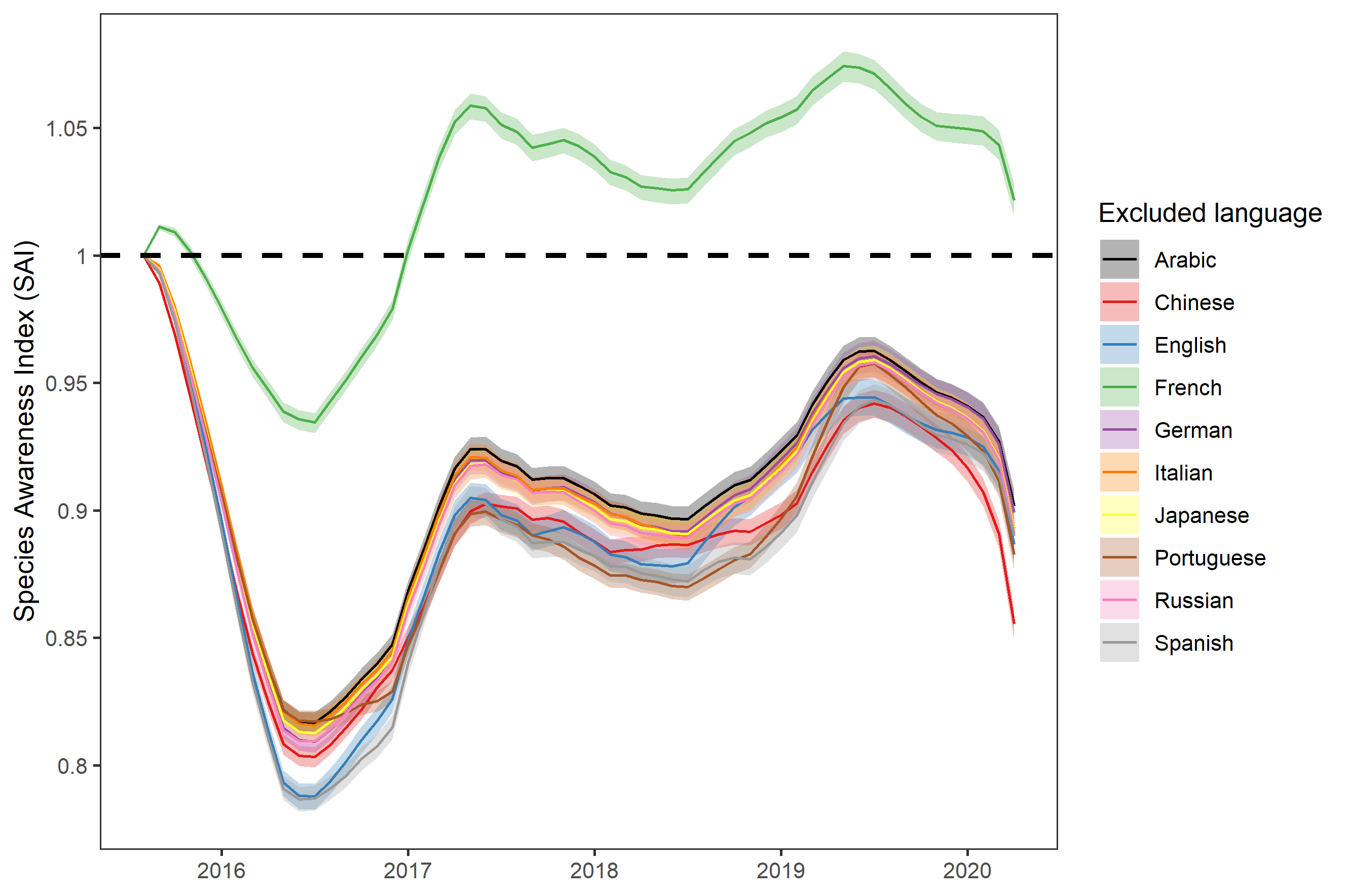

Figure S6. The overall SAI for all species pooled, jack-knifed for language. Each line represents the mean of the bootstrapped indices, and each band the 2.5^th^ and 97.5^th^ percentiles The colour of each line represents the overall SAI if that given language is excluded, providing an indication as to how single languages influence the overall SAI: black (Arabic), red (Chinese), blue (English), green (French), purple (German), orange (Italian), yellow (Japanese), brown (Portuguese), pink (Russian), and grey (Spanish). Here the French language has a significant effect on the overall trend, meaning it was removed in the main text.

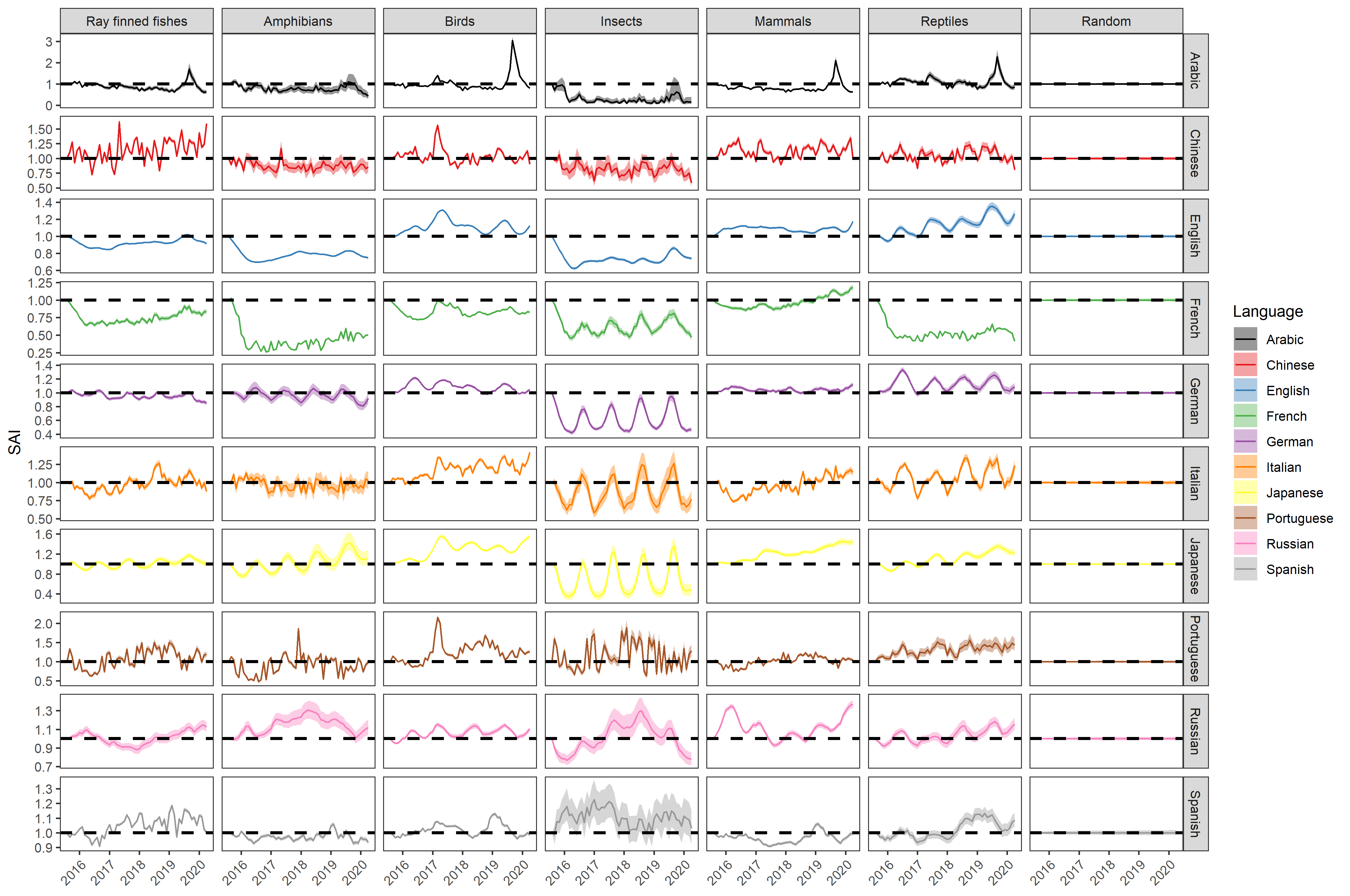

Figure S7. The overall SAI for each taxonomic class in each language before a final loess smooth (span = 0.3) is applied to each species page SAI. More tortuous trends are those for which the views for a given page tend to be lower. The “Random” column represents the random index adjusted for itself, indicating that the adjustment is functioning correctly.

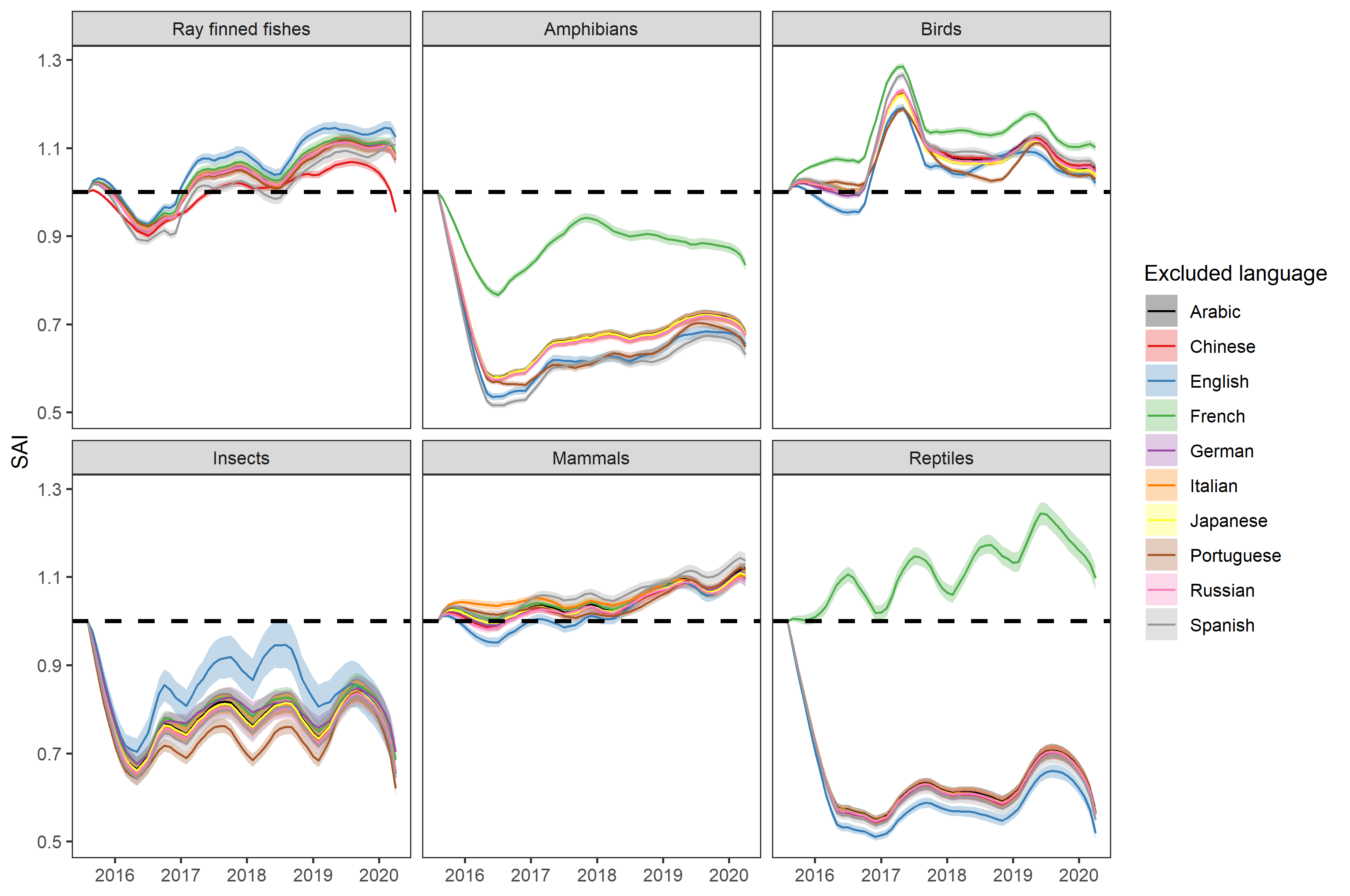

Figure S8. The overall SAI for each taxonomic class, jack-knifed for language. Each line represents the mean of the bootstrapped indices, and each band the 2.5^th^ and 97.5^th^ percentiles. The colour of each line represents the overall SAI if that given language is excluded, providing an indication as to how single languages influence the overall SAI: black (Arabic), red (Chinese), blue (English), green (French), purple (German), orange (Italian), yellow (Japanese), brown (Portuguese), pink (Russian), and grey (Spanish). Here the French language has a significant effect on the overall trend, meaning it was removed in the main text.

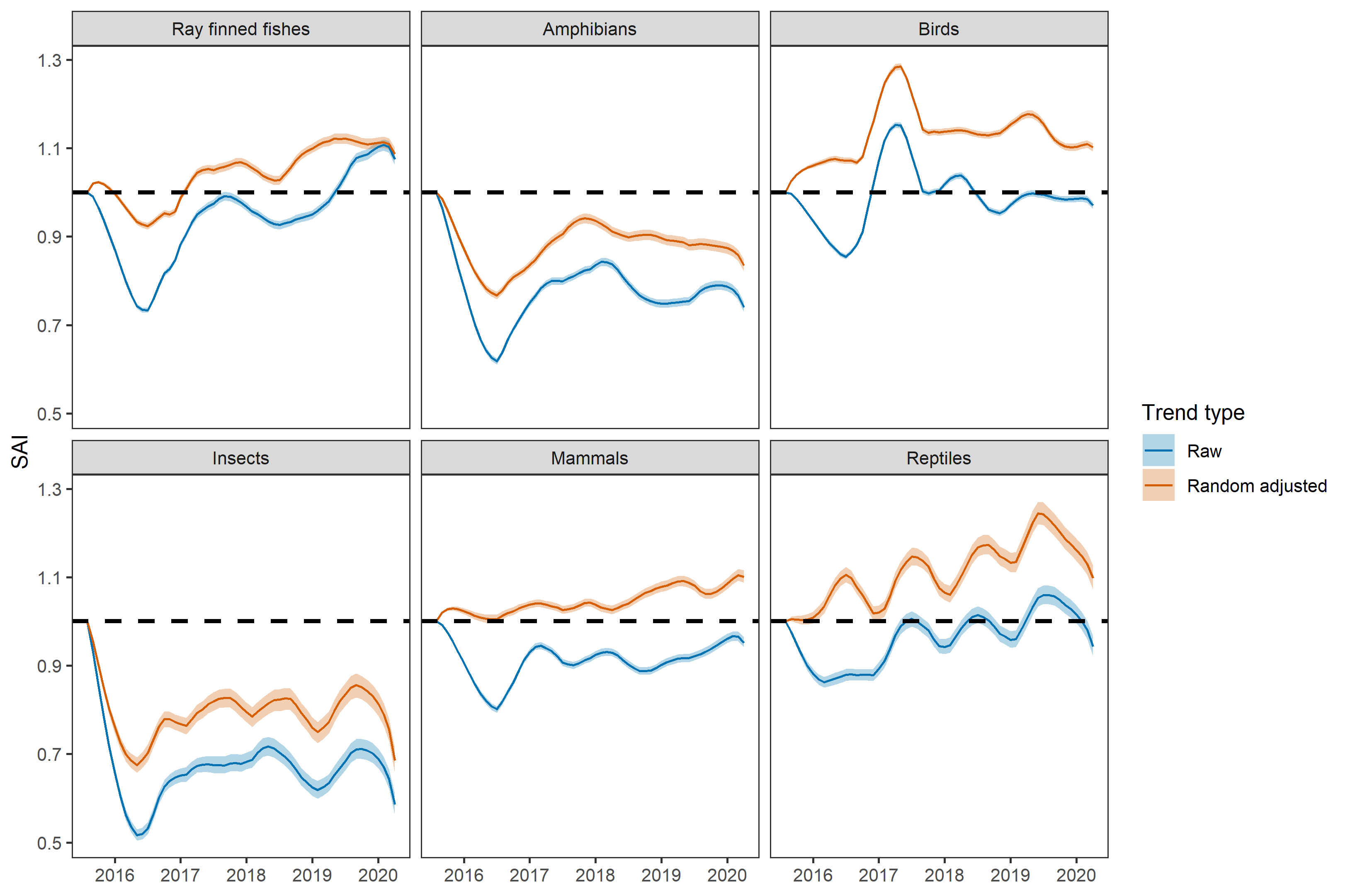

Figure S9. The overall SAI and SAI without a random adjustement for each taxonomic class. Each line represents the mean of the bootstrapped indices, and each band the 2.5^th^ and 97.5^th^ percentiles. Blue lines represent the trend without subtracting the random index, and orange lines represent the trend with the random index subtracted. In other words, the orange line represents the overall trend in awareness without the underlying residual trend of Wikipedia.

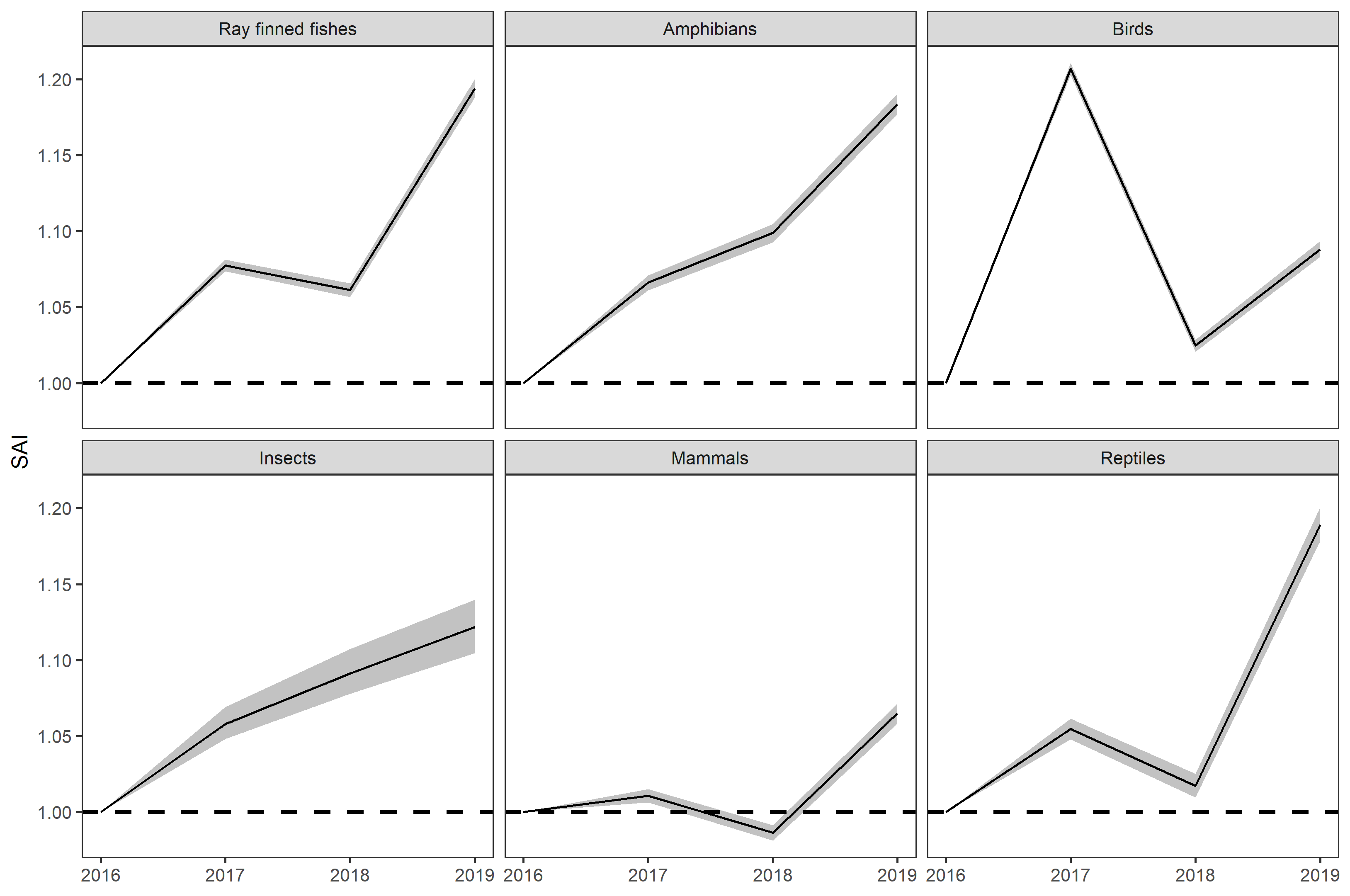

Figure S10. The overall SAI for each taxonomic class, based on the average monthly views per year. Here all languages (Arabic, Chinese, English, French, German, Italian, Japanese, Portuguese, Russian, are Spanish) are included, with no additional loess smoothing of the species page SAI. Although annual trends may be more useful in the future, given this current time series is represented by only four points we opted not to include the annual trends in the main text.

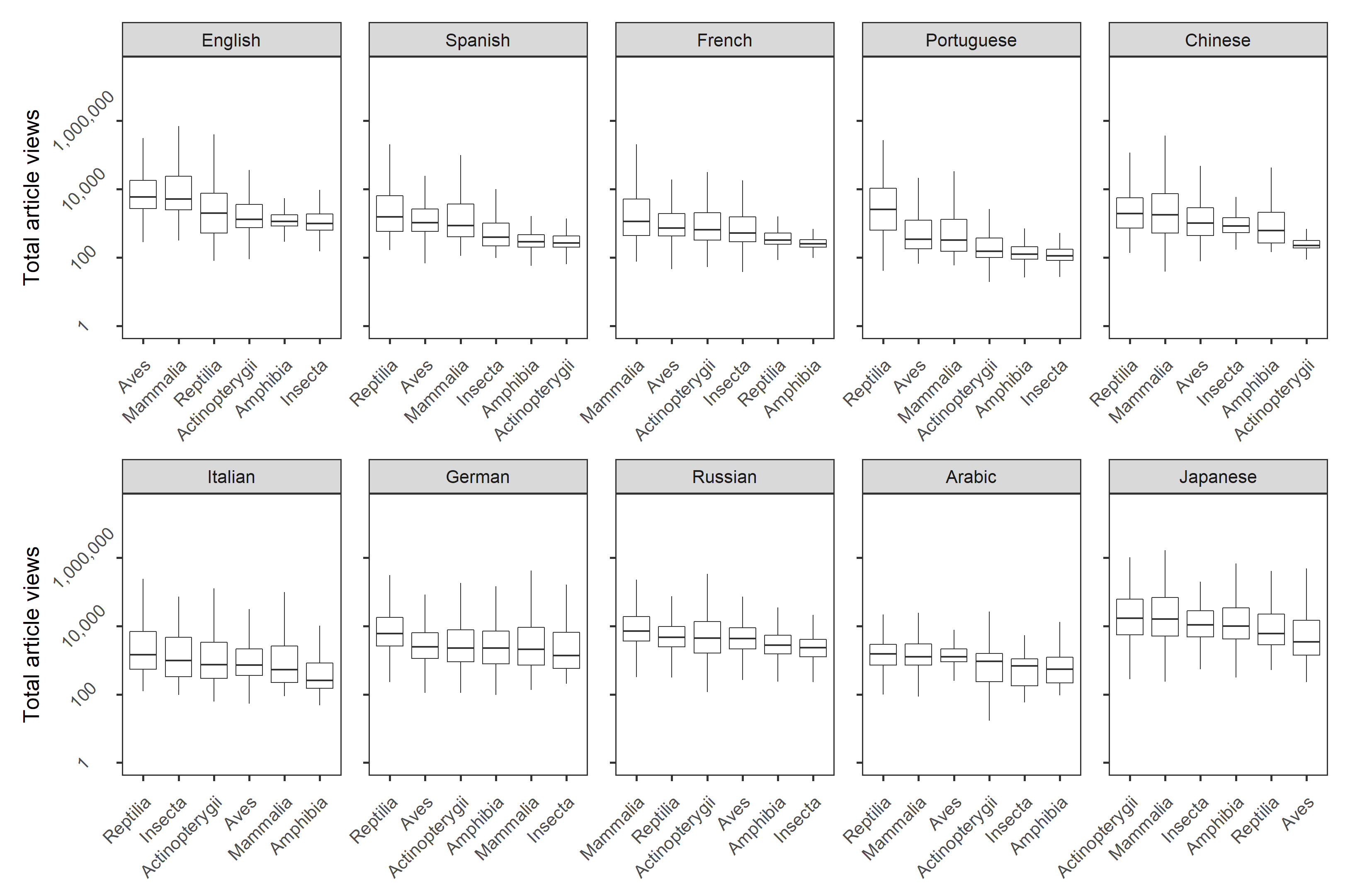

Figure S11. The distribution of page views for each taxonomic class in each language over the period 1^st^ July 2015- 31^st^ March 2020, as an indication of absolute awareness.

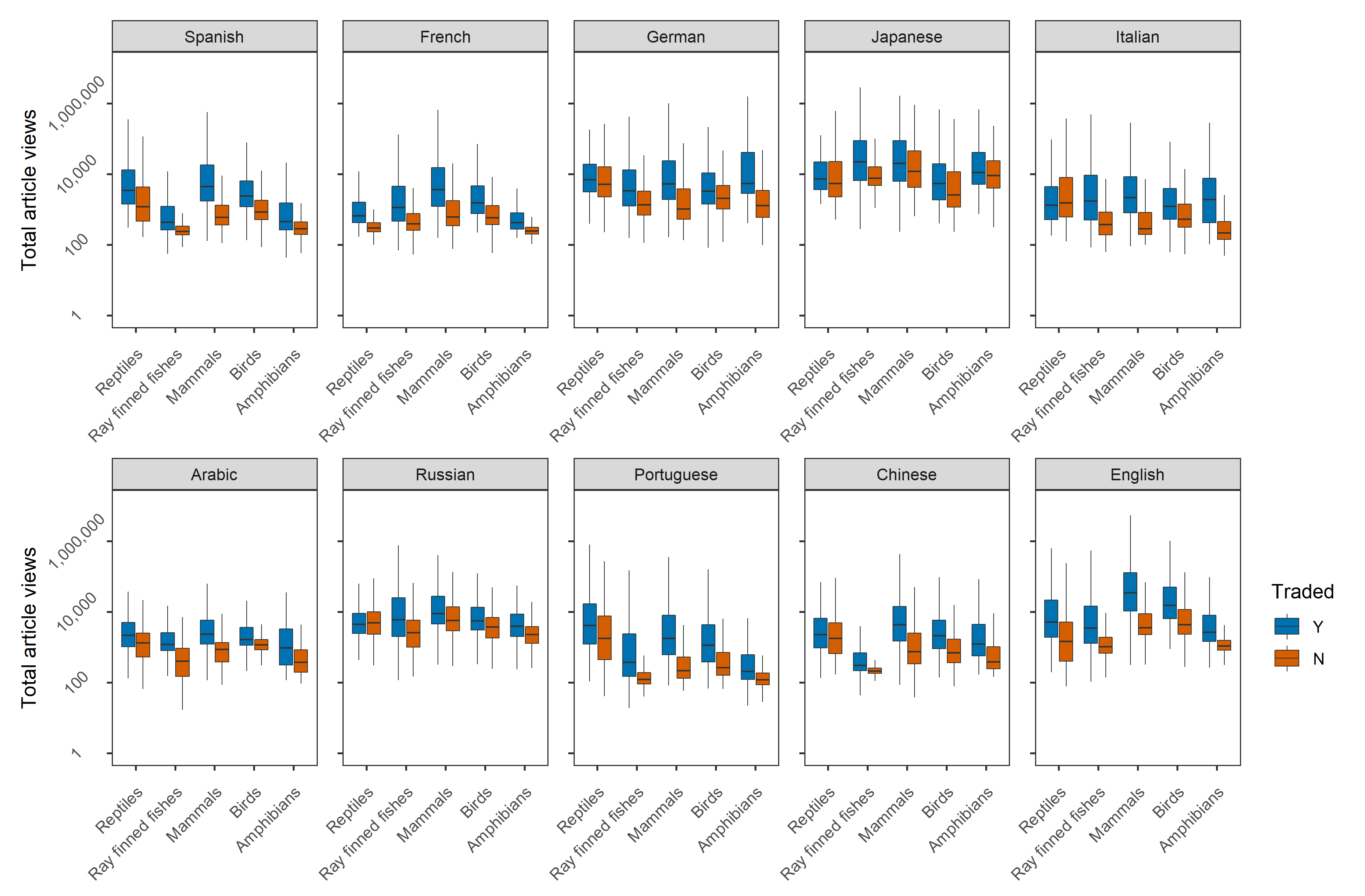

Figure S12. The distribution of views for traded and non-traded species in each taxonomic class in each language, for the period 1^st^ July 2015- 31^st^ March 2020. Blue boxplots represent species that are known to be traded, and red boxplots represent species that are not known to be traded.

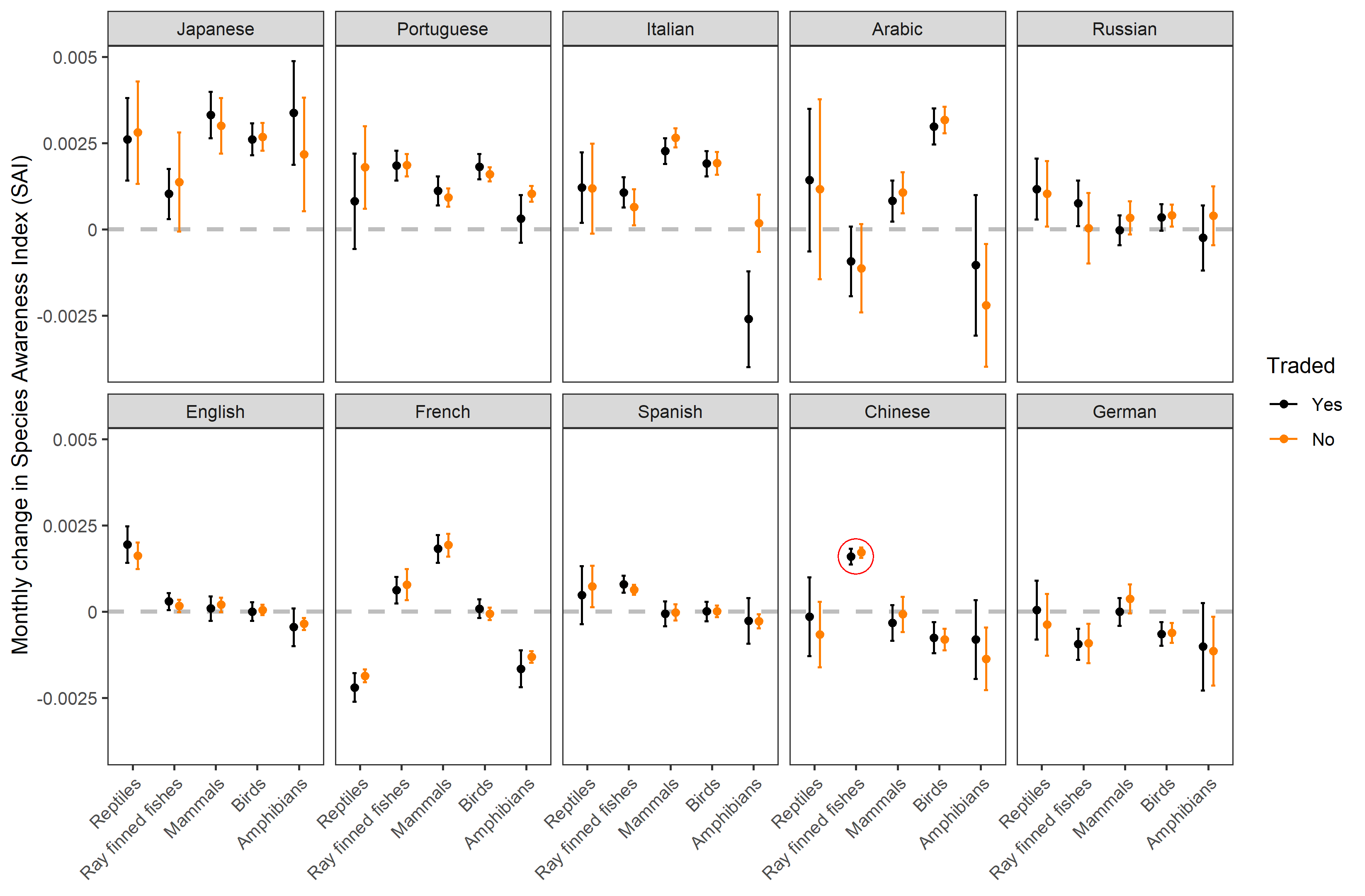

Figure S13. Average monthly rate of change for the species page SAI for 6 taxonomic classes across 10 Wikipedia languages. Errors bars represent the predicted values of a linear model, fitting average monthly change in the species page SAI as a function of taxonomic class, Wikipedia language, trade status (Y/N), and their interaction. Black bars refer to species that are traded, whereas orange bars refer to species which are not traded. The red circle highlights the Chinese ray-finned fish, in which both traded and non-traded species have a similar rate of change (see Discussion).

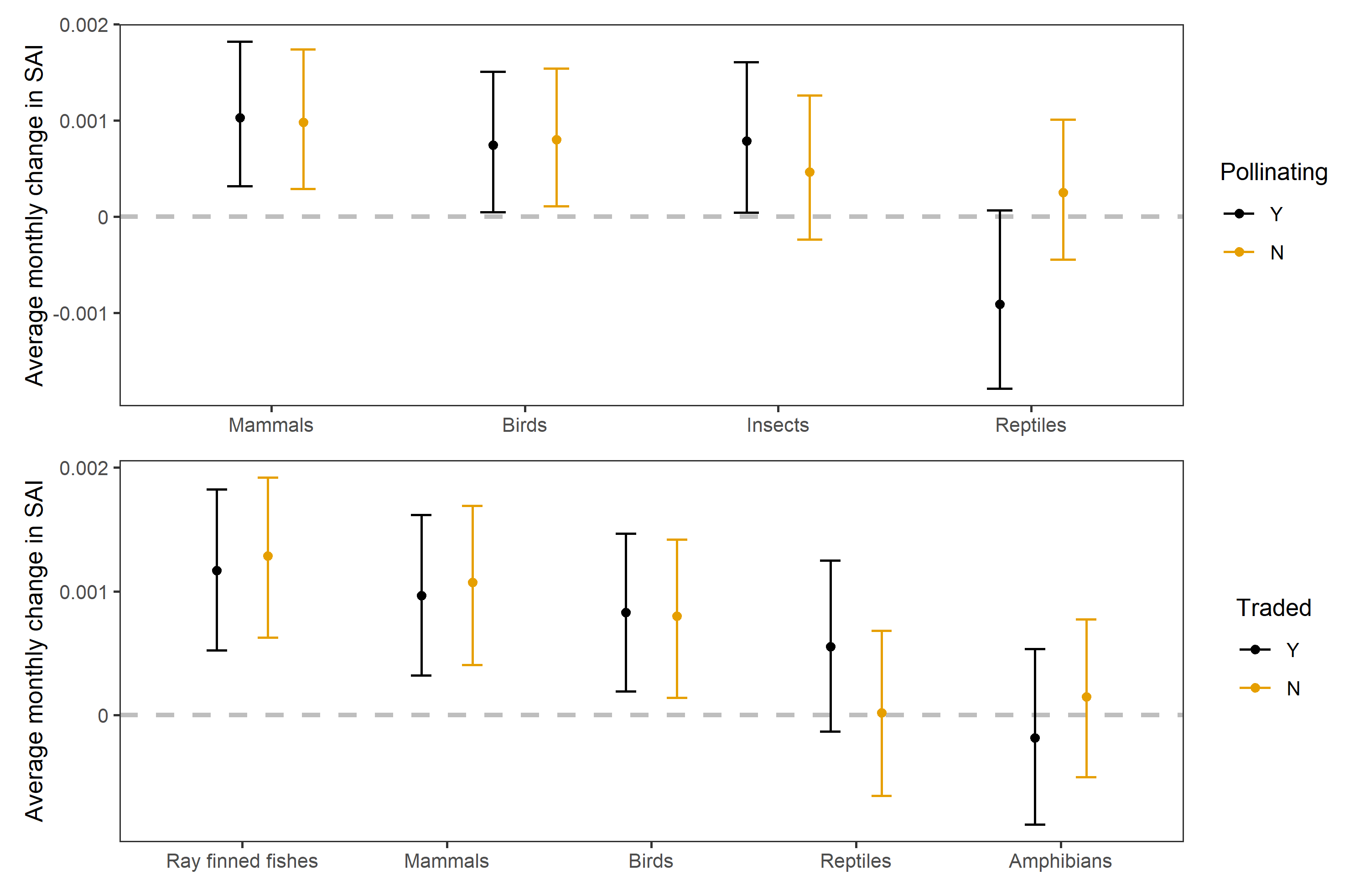

**Figure S14. Average monthly rate of change for the species page SAI as a function of pollination contribution and trade. Top: a mixed effects model for average rate of change in species page SAI as a function of taxonomic class, pollination contribution (Y/N), their interaction, and language (random effect). Bottom: a mixed effects model for average rate of change in species page SAI as a function of taxonomic class, traded species, and language (random effect). Effect sizes were calculated for both panels by drawing fixed effects 1,000 times based on the variance-covariance matrix, and then calculating the median value (shown as points), and 2.5^th^ and 97.5^th^ percentiles (shown as error bars).** **Note that monthly rate of change for reptiles is much lower here given the inclusion of the French Wikipedia (see jack-knife figures S6 and S8).**

**Table S1. ANOVA table for a mixed effects linear model predicting the log10(total views) for each species page as a function of taxonomic class, pollination contribution, their interaction, and a random effect for language.**

| **Fixed effect** | **Sum of squares** | **Mean square** | **F value** | **P value** |
| --- | --- | --- | --- | --- |
| Pollination contribution | 0.18 | 0.178 | 0.3869 | 0.5339 |
| Taxonomic class | 889.74 | 296.579 | 645.8999 | <0.001 |
| Pollination contribution * Taxonomic class | 78.57 | 26.192 | 57.0409 | <0.001 |

**Table S2. ANOVA table for a mixed effects linear model predicting the log10(total views) for each species page as a function of taxonomic class, trade status, their interaction, and a random effect for language.**

| **Fixed effect** | **Sum of squares** | **Mean square** | **F value** | **P value** |
| --- | --- | --- | --- | --- |
| Trade status | 5212.6 | 5212.6 | 15206.44 | <0.001 |
| Taxonomic class | 5813.2 | 1453.3 | 4239.59 | <0.001 |
| Trade status * Taxonomic class | 401.5 | 100.4 | 292.78 | <0.001 |

**Table S3. ANOVA table for a mixed effects linear model predicting the average rate of change in species page SAI as a function of taxonomic class, pollination contribution, their interaction, and a random effect for language.**

| **Fixed effect** | **Sum of squares** | **Mean square** | **F value** | **P value** |
| --- | --- | --- | --- | --- |
| Pollination contribution | 0.00013237 | 0.00013237 | 4.3662 | 0.036662 |
| Taxonomic class | 0.00198949 | 0.00066316 | 21.8750 | <0.001 |
| Pollination contribution * Taxonomic class | 0.00047564 | 0.00015855 | 5.2299 | 0.001314 |

**Table S4. ANOVA table for a mixed effects linear model predicting the average rate of change in species page SAI as a function of taxonomic class, trade status, their interaction, and a random effect for language.**

| **Fixed effect** | **Sum of squares** | **Mean square** | **F value** | **P value** |
| --- | --- | --- | --- | --- |
| Trade status | 0.0000017 | 0.00000165 | 0.0422 | 0.8373260 |
| Taxonomic class | 0.0118755 | 0.00296888 | 75.6620 | <0.001 |
| Trade status * Taxonomic class | 0.0008680 | 0.00021699 | 5.5301 | <0.001 |

**Table S5. ANOVA table for a linear model predicting the average rate of change in species page SAI as a function of taxonomic class, language, and their interaction.**

| **Fixed effect** | **Degrees of freedom** | **Sum of squares** | **Mean square** | **F value** | **P value** |
| --- | --- | --- | --- | --- | --- |
| Taxonomic class | 5 | 0.0321 | 0.0064104 | 162.044 | < 0.001 |
| Language | 9 | 0.0631 | 0.0070092 | 177.181 | < 0.001 |
| Taxonomic class * Language | 45 | 0.0599 | 0.0013317 | 33.662 | < 0.001 |

**Table S6. AIC and ΔAIC values for a set of mixed effects generalised linear models: one fitting log10(total views) as a function of an interaction between pollination contribution and taxonomic class, and three a series of candidate null models. All models were fit with one random effect (language).**

| **Model fixed effects** | **AIC** | **ΔAIC** |
| --- | --- | --- |
| Pollination contribution * Taxonomic class | 183919.7 | 0 |
| Pollination contribution | 188112.0 | 4192.25 |
| Taxonomic class | 184245.5 | 325.76 |
| Intercept (1) | 188306.3 | 4386.6 |

**Table S7.** **AIC and ΔAIC values for a set of mixed effects generalised linear models: one fitting log10(total views) as a function of an interaction between trade status and taxonomic class, and three a series of candidate null models. All models were fit with one random effect (language).**

| **Model fixed effects** | **AIC** | **ΔAIC** |
| --- | --- | --- |
| Trade status * Taxonomic class | 254981.1 | 0 |
| Trade status | 276230.9 | 2124.98 |
| Taxonomic class | 276276.0 | 2129.48 |
| Intercept (1) | 295182.9 | 4020.18 |

**Table S8. AIC and ΔAIC values for a set of mixed effects generalised linear models: one fitting the average monthly rate of change in species page SAI as a function of an interaction between pollination contribution and taxonomic class, and three a series of candidate null models. All models were fit with one random effect (language).**

| **Model fixed effects** | **AIC** | **ΔAIC** |
| --- | --- | --- |
| Pollination contribution * Taxonomic class | -583810.4 | 0 |
| Pollination contribution | -583776.7 | 33.7 |
| Taxonomic class | -583865.6 | -55.2 |
| Intercept (1) | -583796.0 | 14.4 |

**Table S9. AIC and ΔAIC values for a set of mixed effects generalised linear models: one fitting the average monthly rate of change in species page SAI as a function of an interaction between trade status and taxonomic class, and one candidate null model. Only one candidate null model was fit given the models for taxonomic class only and intercept only were fit to a different number of observation (trade data was not found for Squamate reptiles). All models were fit with one random effect (language).**

| **Model fixed effects** | **AIC** | **ΔAIC** |
| --- | --- | --- |
| Trade status * Taxonomic class | -914369.5 | 0 |
| Trade status | -914065.2 | 304.2876 |

**Table S10 AIC and ΔAIC values for a set of linear models: one fitting the average monthly rate of change in species page SAI as a function of an interaction between taxonomic class and language, and three a series of candidate null models.**

| **Model fixed effects** | **AIC** | **ΔAIC** |
| --- | --- | --- |
| Taxonomic class * Language | -950543.4 | 0 |
| Taxonomic class | -947577.1 | 2966.3 |
| Language | -948712.2 | 1831.2 |
| Intercept (1) | -946797.9 | 3745.5 |
